# Characterizing BOLD activation patterns in the human hippocampus with laminar fMRI

**DOI:** 10.1101/2024.07.04.602065

**Authors:** Viktor Pfaffenrot, Antoine Bouyeure, Carlos Alexandre Gomes, Sriranga Kashyap, Nikolai Axmacher, David G Norris

## Abstract

The human hippocampus has been extensively studied at the macroscale using functional magnetic resonance imaging (fMRI) but the underlying microcircuits at the mesoscale (i.e., at the level of layers) are largely uninvestigated in humans. We target two questions fundamental to hippocampal laminar fMRI: How does the venous bias affect the interpretation of hippocampal laminar responses? And can we establish a benchmark laminar fMRI experiment which robustly elicits single-subject hippocampal activation utilizing the most widely applied GRE-BOLD contrast? We comprehensively characterized GRE-BOLD responses as well as T_2_*, tSNR and physiological noise as a function of cortical depth in individual subfields of the human hippocampus. Our results show that the vascular architecture differs between subfields leading to subfield-specific laminar biases of GRE-BOLD responses. Using an autobiographical memory paradigm, we robustly acquired depth-specific BOLD responses in hippocampal subfields. In the CA1 subregion, our results indicate a more pronounced trisynaptic path input rather than dominant direct inputs from entorhinal cortex during autobiographical memory retrival. Our study provides unique insights into the hippocampus at the mesoscale level, and will help interpreting hippocampal laminar fMRI responses and allow researchers to test mechanistic hypotheses of hippocampal function.

## 1. Introduction

The hippocampus is known to play a vital role in core cognitive functions including learning and memory. It is subdivided into several subfields, including the dentate gyrus (DG), cornu ammonis (CA) 1-4, and the subiculum.

While the human hippocampus has been extensively studied at the macroscale using functional magnetic resonance imaging (fMRI), the underlying microcircuits at the mesoscale (e.g., laminar level) have barely been investigated^1^. In contrast to the six-layered neocortex, the hippocampus contains three (partially subdivided) layers. Furthermore, animal research has shown that in stark difference to the classical canonical microcircuits in the neocortex, differentiating feedforward and feedback pathways^2^, the hippocampal projections are primarily feedforward. Nevertheless, inputs to hippocampal pyramidal cells can be differentiated based on their position along the dendritic tree (a schematic depiction is given in supplementary figure S1): While synaptic connections along the “trisynaptic pathway”^3–5^ (entorhinal cortex (EC) → DG → CA3 → CA1) terminate at perisomatic CA3 and CA1 layers (i.e., St. pyramidale and St. radiatum), direct EC projections terminate at distal apical dendrites (in St. lacunosum-moleculare) of CA1 and CA3; CA3 receives additional input to proximal apical dendrites via its abundant recurrent collaterals^6^.

Recent advances in ultra-high field (UHF) fMRI made it possible to obtain structural ^7^ and physiological^1,8^ information from the human hippocampus with sub-millimeter resolution. This high spatial resolution allows fMRI responses to be probed as a function of depth (also known as laminar fMRI). This methodology holds the key to investigating the above-mentioned projections non-invasively in humans. However, methodological limitations and physiological constraints pose challenges for the design and interpretation of cortical depth-dependent fMRI experiments of the hippocampus. The most-widely used contrast for laminar fMRI is the gradient echo (GRE) blood oxygenation level dependent (BOLD) contrast due to its high sensitivity^9–11^ and efficient sequence implementation with echo planar imaging (EPI) readouts. However, GRE-BOLD responses at the laminar level are weighted toward large venous vessels, the well-known draining vein bias. While the main alternative to GRE-BOLD, vascular space occupancy (VASO), can measure laminar responses free of venous contamination in the neocortex^12–14^, it suffers from an intrinsically low temporal efficiency and reduced sensitivity to CBV changes in inferior brain structures which renders VASO currently unviable in the hippocampus^15^. In the neocortex, the venous drainage pattern and its effect on laminar fMRI has been thoroughly described^16–19^. However, because both the anatomy and the vascularization of the hippocampus are considerably different from those in to the neocortex, it is currently not known how the venous bias would affect hippocampal laminar BOLD responses. Hence, it is imperative to characterize the distribution of large blood vessels and their influence on hippocampal laminar fMRI. Furthermore, even though UHF fMRI provides a higher baseline signal-to-noise ratio (SNR) and an improved BOLD contrast^20,21^, it is also more prone to artifacts associated with main magnetic field (B_0_) and transmission field (B_1_) inhomogeneities^22,23^. The anatomical location of the hippocampus renders it particularly susceptible to these inhomogeneities and hence a challenging region for imaging. Therefore, it is necessary to characterize high-resolution functional imaging of the human hippocampus in terms of depth-dependent mapping of the T_2_* relaxation time, signal stability and noise characteristics. Only with this information, we can interpret high-resolution GRE-BOLD activation patterns.

As a prerequisite for transferring neuroscientific fMRI experiments from low to high spatial resolutions and for validating other fMRI contrasts and/or sequences, a benchmark experiment is necessary. In the context of autobiographical memory (AM), neuropsychological and fMRI studies have shown that several subfields of the hippocampus are involved in AM. The subfields include CA1^24^, CA3^25–27^ and the subiculum^28,29^, making an AM experiment an ideal candidate for benchmarking.

In this work, we leverage sub-millimeter GRE-BOLD fMRI performed at 7 T to answer two fundamental questions on laminar fMRI of the human hippocampus:

1) What is the laminar distribution of venous draining patterns in different hippocampal subregions and how does it affect the interpretation of hippocampal laminar responses?
2) Can we perform a benchmark laminar fMRI experiment which robustly elicits single-subject hippocampal activation using the GRE-BOLD contrast? Can we obtain interpretable results despite the venous bias?

To answer the first question, we invited nine participants to perform a breath-holding experiment while acquiring high-resolution multi-echo T_2_*-weighted images. Additionally, we acquired high-resolution structural T_1_- and susceptibility-weighted images as well as time-of-flight (TOF) images. The hypercapnic challenge led to a ubiquitous increase in blood flow and thus to a widespread BOLD response. Sampling at different echo times allowed us to differentiate signal changes inside venous vessels from signal changes in the parenchyma. To address the second aim, the same cohort was scanned while performing an autobiographical memory task on a separate day. This task was shown to elicit reliable single subject activity at high-spatial resolution using GRE-BOLD^29^. By modeling the hippocampus as a folded surface, we investigated unique depth-dependent features of the hippocampal subfields during memory processing contrasted against a control condition, and during the construction (i.e., initial search phase for a specific memory) vs. the elaboration (i.e., reliving the perceptual details of the memory with autonoetic consciousness) of autobiographical memories^30^. We linked the results of the two experiments to interpret the resulting profiles during AM with regard to neural activity and to obtain MRI parameters relevant for future neuroimaging studies of the hippocampus. Altogether, the presented methods and results will facilitate interpretation of hippocampal laminar fMRI responses and validation of other fMRI contrasts. Furthermore, our findings provide mechanistic insights into hippocampal function at the mesoscale.

## 2. Methods

All data were acquired on a 7 T system (MAGNETOM Terra, Siemens Healthcare, Erlangen, Germany) equipped with a 1 channel transmit, 32 channel receive head coil (NOVA Medical Inc.). Nine healthy participants (3 female, 26.4 ± 6.0 years) were scanned in both experiments after giving informed consent according to the guidelines of the local ethics committee. All processing and analysis code can be found in the public github repository (https://github.com/viktor-pfaffenrot/hippocampus_laminarfMRI_code).

### 2.1 Breath-holding experiment

#### 2.1.1 MR Data Acquisition

##### 2.1.1.1 Anatomical Scans

Whole brain T_1_-weighted anatomical reference data were acquired with an MP2RAGE sequence^31^ with fat navigators^32^ at 0.75 mm isotropic resolution. To obtain veno- and angiograms, vendor-provided susceptibility-weighted (SWI) and time-of-flight (TOF) sequences were used to acquire images at 0.3 x 0.3 x 0.6 mm resolution (interpolated to 0.15 mm in-plane). The field-of-view (FOV) was positioned such that the long axis of the FOV was parallel to the long axis of the hippocampus (see supplementary figure S2A). Sequence details are shown in table 1.

**Table 1:**
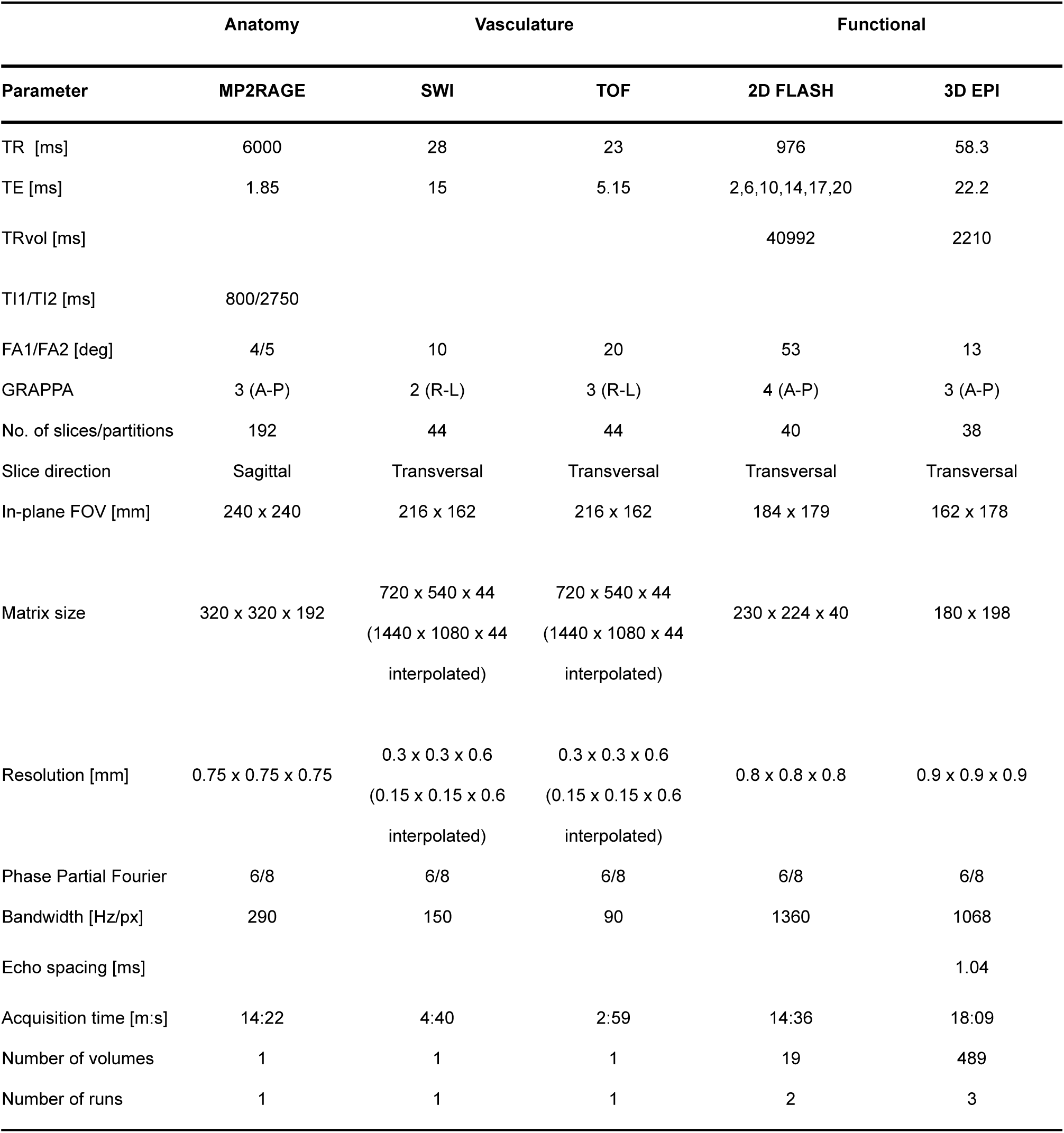
MRI acquisition parameters.

##### 2.1.1.2 Breath-hold Scans

For the breath-hold experiment, data were acquired at 0.8 mm isotropic resolution using a 2D multi-echo FLASH sequence. The FOV was positioned similarity to that of the SWI and TOF acquisitions. Echo times were TE = 2, 6, 10, 14, 17, 21 ms. An additional phase stabilization echo was acquired at the start of each echo train. The effect the phase stabilization has on the data quality is shown in supplementary figure S2B-E. Shot-to-shot TR was 976 ms and 40 slices were acquired resulting in a volume TR of TR_vol_ = 40.992 ms. For each participant, two runs of 19 volumes were acquired with each run lasting 14:36 min.

##### 2.1.1.3 Breath-hold Paradigm

The maximum BOLD response depends on the breath-hold duration and, for the volume TR used in this study, peaks about 10 seconds after the participant started to breathe again^33^. To account for this delay, the paradigm was designed as follows: One volume of normocapnia (labeled as ‘rest’) was followed by a transition volume where the participant was cued to hold their breath in 5 seconds. The participants were instructed to slowly inhale into their abdomen once the breath-holding command appeared and to hold their breath until the command disappeared. Ten seconds before the breath-hold period ended, the acquisition of the active volume started. The temporal shift between paradigm and image acquisition was designed such that the peak of the BOLD response (10 seconds after hypercapnia) coincided with the acquisition of the k-space center of the ‘active’ volume. The participants were instructed to slowly exhale after the breath-hold command disappeared. The block of ‘rest, ’transition’, ’active’ was repeated 6 times and a run ended with a rest volume.

#### 2.1.2 Data Processing

##### 2.1.2.1 Anatomical Scans

MP2RAGE UNI and INV1 images were pre-processed using *presurfer*^34^. Both processed images were denoised using the BM4D filter^7,35^. The image quality improvement is shown in supplementary figure S3A,B and filter parameters deviating from default can be found in our github repository. To obtain surface boundaries and hippocampal subfield segmentations, the snakemake^36^ implementation of hippunfold^37,38^ (https://hippunfold.readthedocs.io.) was used. In particular, the denoised INV1 image was used as a proxy for a T_2_-weighted input due its T_2_-like image contrast (see supplementary figure S3C-E).

Both SWI and TOF images were denoised using BM4D with a fixed noise estimate σ of 30. SWI was receiver bias-field corrected and filtered for veins using a Frangi vesselness filter^39^ (α = β = 0.5, Fig. 1B). TOF images were filtered for arteries using a custom filter based on a Haar 2-D wavelet transform. Both vessel-filtered images were subsequently registered to the T_1_-weighted image using ITK-SNAP Version 3.6.0^40^ and SPM12 Version 7487 (Welcome Trust Centre for Neuroimaging, London). Finally, the resampled images were projected onto the inner and outer surface of the hippocampus as given by hippunfold using the connectome workbench’s^41^ command *volume-to-surface-mapping*.

**Fig. 1:**
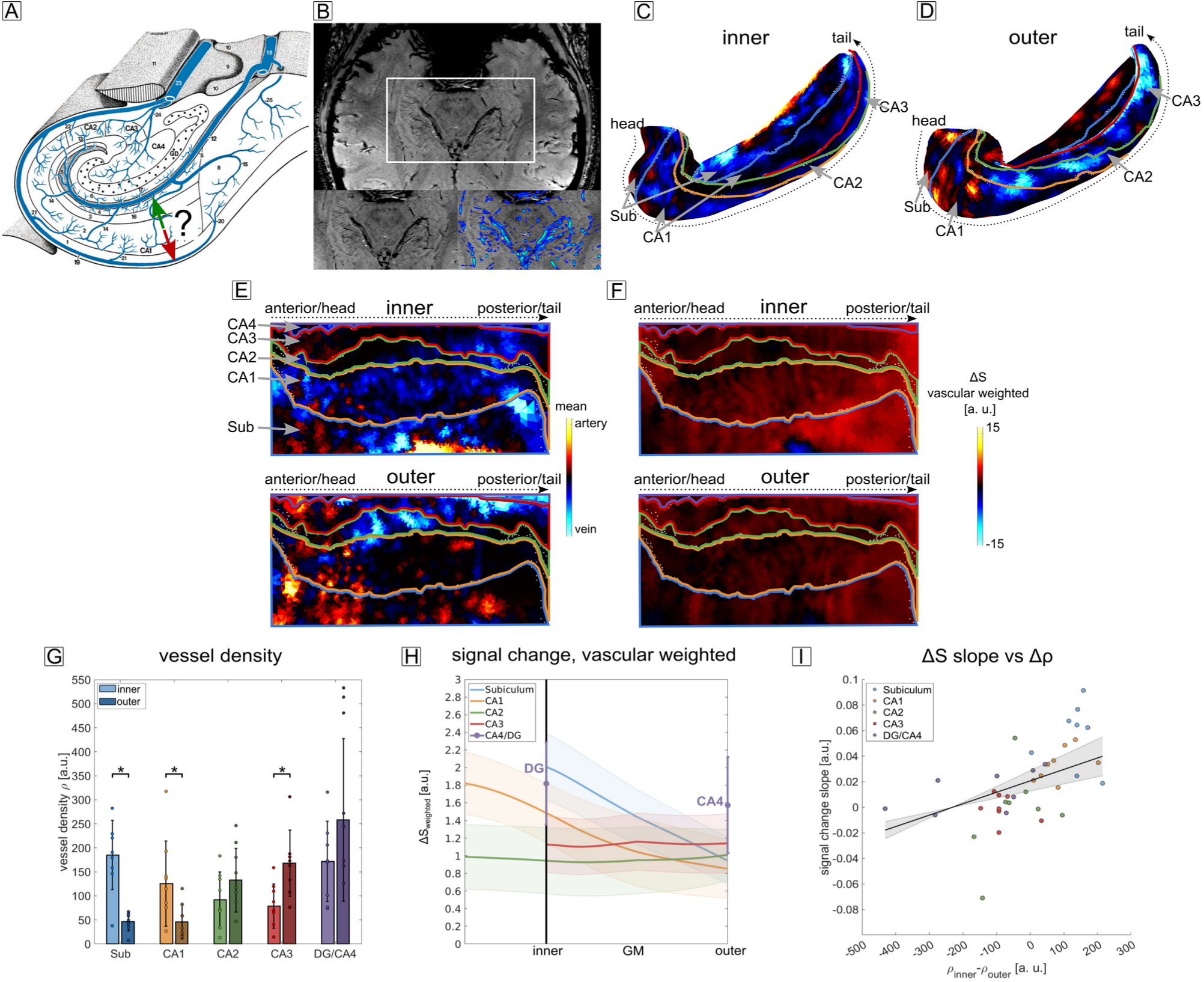
Vascular distribution and beath-hold induced signal change. **A** Schematic of the venous vessel distribution in the human hippocampus. The hippocampus exhibits two main venous drainage pathways: One along the sulcal intrahippocampal veins (green arrow) and one along the subependymal intrahippocampal veins (red arrow). From a laminar fMRI perspective, it is unclear in which direction the venous bias present in GRE-BOLD points to. E.g. for CA1, the signal change could either be biased toward the inner surface (in direction of the green arrow) or toward the outer surface (in direction of the red arrow) (figure adapted from Duvernoy et al.^26^). **B** High-resolution SWI image filtered for veins (blue structures in zoomed section). **C-D** Subject-averaged vascular distribution of large vessels at the inner (**C**) and outer (**D**) surface (veins and arteries in blue and red, respectively) shown on the folded hippocampus. Colored borders represent the demarcation between subfields. The venous density exhibits a clear subfield-specific differentiation between the inner and outer surface. **E** Same data as in **C, D** plotted on the unfolded hippocampal surfaces. From bottom to top: Subiculum, CA1-4. **F** Breath-hold induced signal change with vascular weighting. The signal changes closely resembles the venous vasculature and is higher at the inner surface for the subiculum and CA1, while for CA2-4, the opposite is the case, although less in magnitude. **G** Venous vessel density ρ for the inner and outer layer of each subfield. Subiculum and CA1 show a significantly higher vessel density at the inner compared to the outer surface. CA3 shows a significantly higher density at the outer compared to the inner surface. **H** Vascular-weighted, subject- and echo-averaged signal change during breath-hold as a function of cortical depth for each subfield (shaded area corresponds to SEM). Note that in case of CA4/DG, no layer sampling was performed (see section 2.1.2.4). Instead, vertices of the entire region were averaged. **I** Relationship between the slope of the signal change curves in **H** and the difference in vessel density Δρ in **G**. An LME revealed a significant main effect of Δρ.

##### 2.1.2.2 Breath-hold Scans

Breath-hold scans were reconstructed offline using BART (v. 0.8.00)^42^. In particular, ESPIRiT^43^ was used to obtain coil sensitivity maps from the autocalibration signal (ACS) before running an iterative SENSE^44,45^ reconstruction with a regularization factor of 0.005 (*pics -r 0.005*). To account for potentially low SNR, all volumes were denoised using the BM4D filter with the same settings as used for the MP2RAGE.

##### 2.1.2.3 Functional Processing

Denoised FLASH images were pre-processed similar to previous work^46,47^ using MATLAB code based on SPM12. In short, after slice-time correction, data were realigned within each run using a brain mask as weights. Motion between runs was accounted for by realigning a biasfield-corrected volume average of the third TE (10 ms) of each run. Registration to the anatomy was performed using ITK-SNAP and a rigid transformation, guided by a mask which excluded areas of strong signal dropout in the sphenoid sinus and maxillary sinuses. All transformation matrices were concatenated and a single resampling operation using a 4th-order spline was applied.

##### 2.1.2.4 Laminar Profile Extraction

The functional data were sampled as a function of cortical depth using custom MATLAB code (https://github.com/viktor-pfaffenrot/hippocampus_laminarfMRI_code/blob/main/ layerfication/VPF_create_hippocampus_layers.m). For each hippocampal subfield, signals were sampled between the surface vertices of the inner and outer hippocampal boundary (see supplementary figure S1 and S4A,B) using 20 equidistant bins. Here, the outer surface refers to the transition region between stratum oriens and parts of St. pyramidale closest to St. oriens. The inner surface corresponds to the transition between St. radiatum and parts of St. pyramidale closest to St. radiatum. To account for the high curvature of the hippocampus, the midthickness depth, provided by hippunfold and obeying the equivolume principle, was used as a landmark in the sampling process. The effect on the laminar profiles is shown in supplementary figure S4C-F. In case of CA1 and CA2, the sampling was extended by 10 bins beyond the inner surface to also sample parts of the St. lacunosum and St. moleculare, i.e. the innermost layers corresponding to the location of distal apical dendrites. In case of CA4 and the DG, only two bins were defined as there is no clear definition of ‘inner’ and ‘outer’ surface. Hence, given the anatomical structure as shown in Fig. 1A (mod. From Duvernoy et al.^48^ Fig. 3.7) and the segmentation of hippunfold, the entire DG was labeled ‘inner’ and the entire CA4 was labeled ‘outer’.

#### 2.1.3 Data Analysis

The projected veno- and angiograms were rescaled to be within the interval [-100,0] and [0,100], respectively, and vessel density was estimated by dividing the number of vertices passing a threshold over the total number of vertices within each subfield. The threshold was -3 a. u. and 1 a .u. for veins and arteries, respectively, and was chosen to reduce the number of falsely classified vessels.

The laminar data of one subject were disregarded due to too excessive motion. For the remaining eight subjects, data of each run were high-pass filtered with a cutoff period of 1/(246 s) and the first and the last volumes of each run were removed to account for edge effects at the boundaries of each run’s time series as a result of high-pass filtering ^49^. Finally, the profiles were averaged over runs and over the rest and the active condition, respectively.

For the rest condition, cortical depth-dependent T_2_* was estimated by fitting a mono-exponential signal decay function to each bin and subfield using MATLAB’s Levenberg-Marquardt nonlinear least squares algorithm.

Breath-hold induced signal change was calculated as breath-hold – normal breathing. To relate signal change to venous vessel density, we averaged over echoes and calculated the slope between outer and inner surface. Although phase stabilization accounts for a high degree of unwanted artifacts, some nuisance signal fluctuations, scaling with echo time, were still present. To reduce their effect, we performed a weighted echo average where the weighting factor penalizes the longer TE and is given by

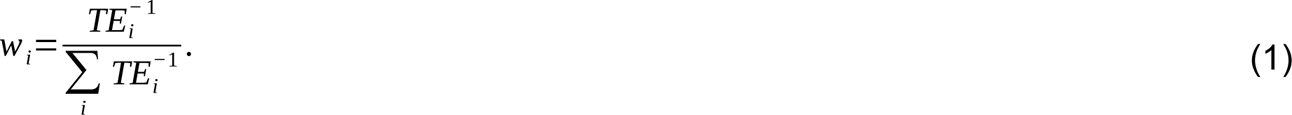

#### 2.1.4 Statistical Analysis

The venous vessel density was compared between the inner and outer surface for each subfield using MATLAB and its build-in Wilcoxon signed-rank test corrected for multiple comparisons with Hochberg’s step-up Bonferroni procedure^50^.

The relationship between the slope of the signal change as a measure for the bias in laminar fMRI, and the vessel density difference was modeled using a linear mixed-effects model in R (version 4.3.3) utilizing the *lme4* package. To investigate whether the difference in vessel density was a significant main effect, we compared a model with the density change as predictor with one including only the random effects. These were modeled as random intercepts for subject and subfield. Both models were compared using an approximated F-test based on the Kenward-Roger approach for small sample sizes ^51^ as implemented in the *pbkrtest* package. Marginal and Conditional R^2^ were obtained with the *performance*^52^ package.

### 2.2 Autobiographical Memory Experiment

#### 2.2.1 MR Data Acquisition

The same participants who performed the breath-hold task were invited for the autobiographical memories (AM) task, hence anatomical data were not re-acquired . Functional AM data were obtained using the GRE-BOLD contrast and a 3D-EPI readout^53^ at 0.9 mm isotropic resolution. The key parameters were: TE/TR/TR_vol_ = 22.2/58.3/2210 ms, FOV = 162x178x34 mm. The remaining sequence details are given in table 1. For each participant, three runs of 489 volumes were acquired with each run lasting 18:09 min. In order to investigate data reliability, two subjects were scanned twice on different days. For distortion correction, the same sequence with reversed-phase encoding was used and 13 volumes were acquired.

##### 2.2.1.1 Autobiographical Memory Paradigm

The functional task was an autobiographical memory paradigm adapted from Leelaarporn et al.^29^ ( see also McCormick et al.^54^). In brief, the task consisted of AM trials as a test condition that were randomly interleaved with mental arithmetic (math) trials as control. In the AM trials, participants were asked to retrieve autobiographical episodic memories following the presentation of general cues (e.g. ’party’, ’pet’, etc.). They had to press a button to indicate when they had found a memory related to the cue, and to vividly imagine the event with autonoetic consciousness with as much detail as possible for the remainder of the trial. The before and after button press phases of each trial thus correspond to the construction and elaboration phases of autobiographical memory retrieval, respectively^30^. Participants were additionally asked to retrieve recent autobiographical memories (no older than two years) to ensure hippocampal activation as remote autobiographical memories have been suggested to evoke less hippocampal activity compared to recent ones^55^.

The AM trials were randomly interleaved with math trials in which subjects had to perform a simple arithmetic operation (e.g. 30+15). Participants were instructed to press a button once they found the result, and to recursively add 3 to the result (e.g., 45+3+3+3…) until the end of the trial in order to keep them engaged for the whole trial duration.

Each AM and mental arithmetic trial lasted 17.6 seconds. The inter-trial interval was a fixation cross with a duration of either 2.2 or 4.4 seconds (in order to be a multiple TR_vol_), which was randomly determined. Each run consisted of 44 trials. Unique cue words and arithmetic operations were presented during each run.

#### 2.2.2 Data Processing

The first three volumes of all runs were disregarded to allow magnetization to reach a steady state. The remaining, online reconstructed magnitude and phase images were used in an offline partial Fourier reconstruction utilizing the POCS algorithm^56,57^. POCS-reconstructed magnitude data were subsequently processing with a custom pipeline based on Advanced Normalization Tools (ANTs)^58^ (https://github.com/ANTsX/ANTs). In short, both functional and reversed-phase encoded images of each run were subject to rigid-body correction, followed by a rigid alignment of the temporal mean of the functional and the reversed-phase encoded data. For distortion correction, an undistorted template image was calculated using the script *antsMultivariateTemplateConstruction2*^59^ employing ANTs’ symmetric normalization (SyN) algorithm. The distortion-corrected template image of each run was subsequently rigidly registration to the anatomy by using ITK-SNAP and the same masking principle as described in section 2.1.2.3. All transformations and distortion-correction warps were concatenated and only one single resampling operation using Lanczos-windowed sinc interpolation was applied to the unprocessed functional data to minimize the resolution loss due to multiple interpolation steps.

After pre-processing, functional data were sampled as described in section 2.1.2.4 to obtain depth-dependent time courses. In addition to the procedure described above, we sought to reduce the venous bias in GRE-BOLD laminar fMRI by masking out venous vessels. To this end, the projected SWI surface, filtered for venous vessels (see section 2.1.2.1 and Fig. 1B), was sampled and used as a venous mask. In particular, surface vertices exhibiting a value below -3 a.u. were labeled as belonging to a vessel and disregarded.

#### 2.2.3 Data Analysis

##### 2.2.3.1 Data Model

Functional data were analyzed in a two-step procedure using robust weighted least squares^60^ within the GLM framework in SPM12. In a first pass, data were analyzed in voxel space. A single design matrix was used with three regressors of interest: The AM condition, split into construction phase and elaboration phase, and the math condition. The six motion parameters were used as nuisance regressors and the data were highpass filtered with a 1/(128 s) cutoff before fitting the model.

It is well known that physiological noise, i.e. respiration and heartbeat, is a major confounding factor in cortical areas near large vessels and ventricles^61,62^ especially at ultrahigh fields^63^. To account for the pronounced physiological noise at the level of the hippocampus, we applied acompCor^64^. In particular, we used two regions of interest (ROIs). For one region, we created a mask of high residuals as given by the residuals map of the first GLM fit. A residual was classified as high if its value was larger than three times the SD of all residuals. This mask should reflect signal fluctuations associated with respiratory and cardiac cycles. A second ROI was manually drawn on the white matter of the brain stem close to the hippocampus. For the first ROI, the number of nuisance regressors was chosen such that 50 % of the variance was explained. For the second ROI, the number of components was fixed to five. In total, this resulted in approximately 11 to 14 additional acompCor regressors for each run. To increase the predictability of the model, the acompCor regressors were orthogonalized with respect to the motion regressors. After obtaining the acompCor regressors, the model was re-fitted at the voxel level. An evaluation of this procedure can be found in the supplementary material (Fig. S5). Finally, two contrasts were generated, one as the difference between construction and elaboration phase (labeled pre > post) and one as the difference between the average of the two AM phases and the control condition (labeled memory > math).

Prior to applying the same GLM model on depth-dependent signals, the layer time courses were baseline z-transformed. That is, the z-transformation was performed using the mean and the SD calculated from the volumes corresponding to the math condition.

##### 2.2.3.2 tSNR, Weisskoff test and physiological noise

Temporal SNR was calculated by dividing the temporal mean over the temporal SD where only the volumes corresponding to the math condition were included.

To investigate cortical depth-dependent physiological noise variations in the hippocampus, we performed a Weisskoff test in a depth-specific manner. The Weisskoff test^65^ was originally devised as a method to assess scanner stability. Originally, the test was performed by continuously increasing the number of voxels in a spatial average before calculating the temporal SD. If the temporal noise is thermal, i.e. of random nature, the noise should decrease with 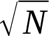, where N is the number of voxels used in the average.

But, at some level of N, any additional voxel in the average does not lead to a noticeable reduction in the noise level. At this point, physiological noise, i.e. structured noise, dominates the overall noise profile. On the laminar level, the Weisskoff test was used to estimate the minimum extend of an ROI that was needed to be in the physiological noise dominated regime^66,67^.

In this work, we averaged over hippocampal hemispheres and performed the Weisskoff test for each subject, each subfield and each depth separately. In particular, we selected profiles at random and varied the number of profiles included in the average from 1, i.e. no averaging, to the maximum number of vertices within each subfield before calculating the temporal noise level (as defined by the temporal SD over time for volumes in the math condition). Finally, we averaged over subjects. This procedure helped us to investigate the laminar variation in physiological noise and to answer the question of whether our current laminar fMRI experiment is performed in the physiological noise dominated regime.

#### 2.2.4 Statistical Analysis

To confirm that the used autobiographical memory task elicits activation in the same areas as previously reported, we conducted a group-level GLM analysis on the memory > math contrast smoothed with a 2.7 mm Gaussian kernel and registered to MNI space. Permutation testing as implemented in the SnPM toolbox^68^ was used to control for the FWE rate. Specifically, 2^9^ permutations were used, and the variance of the model was smoothed with the same kernel as recommended by the developer of the toolbox for small sample sizes. The resulting pseudo t-statistic was thresholded at .05 significance level.

All group-level laminar statistical analyses were performed in R (version 4.3.3) for both functional contrasts (memory > math and pre > post). To assess whether there is a significant main effect of layer, i.e. whether the profiles resemble each other, we performed a linear mixed model comparison using the *lme4* package for each subfield. We compared three models: A baseline model of the form *z ∼ (1|subject),* where *z* is the baseline-transformed z-value and *subject* is the random effect, a linear model (*z ∼ depth + (1| subject)*), where *depth* is the cortical depth as the main effect, and a quadratic model (*z ∼ depth + depth*^2^ *+ (1|subject)*). In particular, we compared the baseline with the linear model and the linear with the quadratic using an approximated F-test based on the Kenward-Roger approach for small sample sizes^51^. This was preferred over other methods, such as the Likelihood Test, as it has been shown to produce acceptable Type 1 Error Rate, particularly for small sample sizes^69^. Multiple comparisons were FDR-corrected using the Benjamini & Hochberg method^70^.

Functional contrasts (memory > math vs. pre > post) were compared by performing a LME analysis for each subfield (except CA4/DG) with depth, contrast and their interaction as main effects (*z ∼ depth + contrast + depth:contrast + (1|subject))* followed by a Type III analysis of variance with Kenward-Roger approximation for degrees of freedom and a FDR correction for multiple comparisons.

To test whether venous vessel masking had an effect on the comparison between cortical depths, we performed a similar analysis as described in the previous paragraph and modeled the response as *z ∼ depth + masking + depth:masking + (1|subject)* followed by testing for significant main and interaction effects for each subfield.

### 2.3 Data Visualization

Results related to the breath-hold experiment (T_2_*, vessel density and breath-hold induced signal change) as well as results related to the autobiographical memory experiment (tSNR) were mapped on the inner and outer surface of the folded and unfolded hippocampus using modified code from the hippunfold toolbox (https://github.com/jordandekraker/hippunfold_toolbox). In addition to the presented visualizations, surface maps and laminar profiles can be interactively explored in our online app (https://viktor-pfaffenrot.shinyapps.io/hippocampus_data_viewer/).

## 3. Results

### 3.1 Venous bias in hippocampal laminar fMRI

To better understand the venous drainage pattern in laminar fMRI of the human hippocampus (Fig. 1A), we mapped the hippocampal vascular anatomy using high-resolution SWI and TOF angiography. By applying filter techniques^39^ (Fig. 1B) followed by mapping on the folded (Fig. 1C, D) and unfolded (Fig. 1E) surface representation of the hippocampus (see supplementary figure S3), we obtained a distinct vascular pattern. Fig. 1C, D shows the subject-averaged vascular distribution of large vessels at the inner (C) and outer (D) surface (veins and arteries in blue and red, respectively). The outer surface putatively corresponds to St. oriens and parts of St. pyramidale closest to St. oriens. The inner surface putatively corresponds to the transition between St. radiatum and parts of St. pyramidale closest to St. radiatum. Fig 1E shows the same data plotted on the unfolded hippocampal surfaces. Comparing between surfaces, Fig. 1C and E show an increased venous vessel density at the inner surface of subiculum and CA1. Both regions show lower densities of large veins at the outer surface (D). In contrast, regions CA2-CA3 (Fig. 1D) exhibit a higher venous density at the outer surface compared to the inner surface. These results appear consistent with the schematic drawings depicted in the Duvernoy atlas (Fig. 1A).

We performed a breath-holding experiment to investigate how these subfield-specific vascular differences translate into biases in the laminar fMRI responses. Because the signal change during breath-holding is completely driven by vascular density distribution (rather than a cognitive task), the signal change distribution should provide information on the expected venous bias. Fig. 1F shows unfolded surface maps of the breath-hold induced, subject- and echo-averaged signal change with vascular weighting, respectively. In the echo average, the vascular weighting penalizes signals acquired at longer echo times which are more contaminated by B_0_-related artifacts not showing any vascular origin (see methods section 2.1.3 for details). The maps of signal change largely resemble the pattern seen in the venous vessel density distribution, i.e. a high signal change at the inner layer of subiculum and CA1 compared to the outer layer and the reverse pattern, although lower in magnitude, for CA2-4.

To quantitatively link venous vessel density ρ to vascular bias, we first statistically compared the differences in vessel density between inner and outer surfaces for each subfield. Fig. 1G shows ρ obtained by aggregating the vertices as shown in Fig. 1E for each subfield. A lighter color represents the inner, a darker color the outer surface, respectively. The venous vessel density is significantly higher at the inner surface for subiculum and CA1 (Wilcoxon signed-rank test W = 36, p_up_ = .031). In case of CA3, the ρ-value is significantly higher at the outer surface (W = 1, p_up_ = .047). CA4/DG shows a trend after multiple comparisons correction (W =3, p_up_ = .078). Fig. 1H show laminar profiles of the subject- and echo-averaged signal change during the breath-hold experiment with vascular-weighting. Single subject and subject-averaged multi-echo laminar profiles can be found in supplementary figure S6. Shaded areas correspond to the SEM across subjects. The profiles show a strong positive slope from the outer toward the inner surface for subiculum and CA1 (blue and orange profiles). CA2 and CA3 (green and red profiles) show a tendency in the opposite direction. The DG exhibits a stronger signal change compared to CA4. Fig. 1I shows the slope of the signal change (corresponding to Fig. 1H) as a function of vessel density difference Δρ computed from Fig. 1G. We conducted a comparison of LME using an approximated F-test (see section 2.2.4). The test revealed a significant effect of vessel density difference (Kenward-Roger approximated F-test, F(1,33.5) = 5.28, p = .028, marginal R^2^ = 0.18, conditional R^2^ = 0.48). The estimated model, plotted as a black line in Fig. 1I (shaded area is the model’s SE), indicates a positive relationship between the slope of the breath-hold induced signal change, i.e. the venous bias, and the vessel density.

In summary, these results show subfield- and layer-dependent differences in venous vasculature and in breath-hold induced signal changes as a measure of physiological bias. They are strongly related and shed light on the venous drainage pattern and how it affects laminar fMRI of the human hippocampus. In particular, subiculum and CA1 show very prominent layer differences in venous drainage, both exhibiting a strong bias from outer to inner layers. CA3 exhibits a bias in the opposite direction, although smaller in magnitude. Layer differences in CA2 and between DG and CA4 appeared less pronounced and did not reach significance after correction for multiple comparisons.

### 2.2 Laminar distribution of autobiographical memory effects

Next, we sought to establish a benchmark laminar fMRI experiment of the human hippocampus utilizing the most-widely used contrast and sequence for laminar fMRI: GRE-BOLD 3D EPI. To this end, inspired by the results of Leelaarporn et al.^29^, we performed an autobiographical memory experiment (see section 2.2.1.1) with the same participants as in the breath-holding experiment.

To verify that the AM experiment elicited similar activation patterns as previously described^29^, we performed a non-parametric group-level analysis at the voxel level (supplementary figure S7) showing activations at the expected locations (primarily anterior body of the hippocampus).

Fig. 2 shows the cortical depth-dependent profiles for the baseline z-transformed and venous vessel masked memory > math contrast for all subjects and subfields along with the subject average (shaded area correspond to SEM). CA4 and DG again were not subdivided into layers. For all remaining subfield, we tested for significant linear and quadratic main effect of depth by performing an approximated F-test with Kenward-Roger approximation for degrees of freedom and restricted maximum likelihood (for details, see section 2.2.4 and Luke^69^). Fig. 2F shows the fitting coefficients obtained (errorbars indicate SE of the coefficient). The asterix indicate whether a higher-order model describes the data statistically better than a lower-order model. The LME-based analysis revealed a significant linear effect of depth in all subfields (F (1,260)> 76, p_FDR_ < .001). A quadratic model fitted the data significantly better than a linear model for the subfields CA1 (F(1,259) = 76, p_FDR_ = 1.5e-15) and CA3 (F(1,178) = 9.4, p_FDR_ = .006). In case of CA3, the quadratic model suggests a higher response In middle layers which can be tentatively interpreted as autobiographical memory being predominantly driven by recurrent collateral inputs to CA3, terminating at proximal apical dendrites, rather than EC inputs to apical distal dendrites or DG inputs to perisomatic layers. CA1 exhibits a prominent peak at inner layers which suggests a dominant input from the trisynaptic pathway, i.e. from CA3, rather than from EC inputs to distal apical dendrites. Obviously, this interpretation should be considered with caution since we could not directly identify anatomical layers and since these anatomical layers show pronounced physiological differences (see Discussion). It is worth noting though that the peak at inner layers of CA1 cannot be explained by the venous bias as our hyperemia experiment would predict a monotonic increase toward St. lacunosum-moleculare (dashed line in Fig. 2B) which is absent in the functional profiles.

**Fig. 2:**
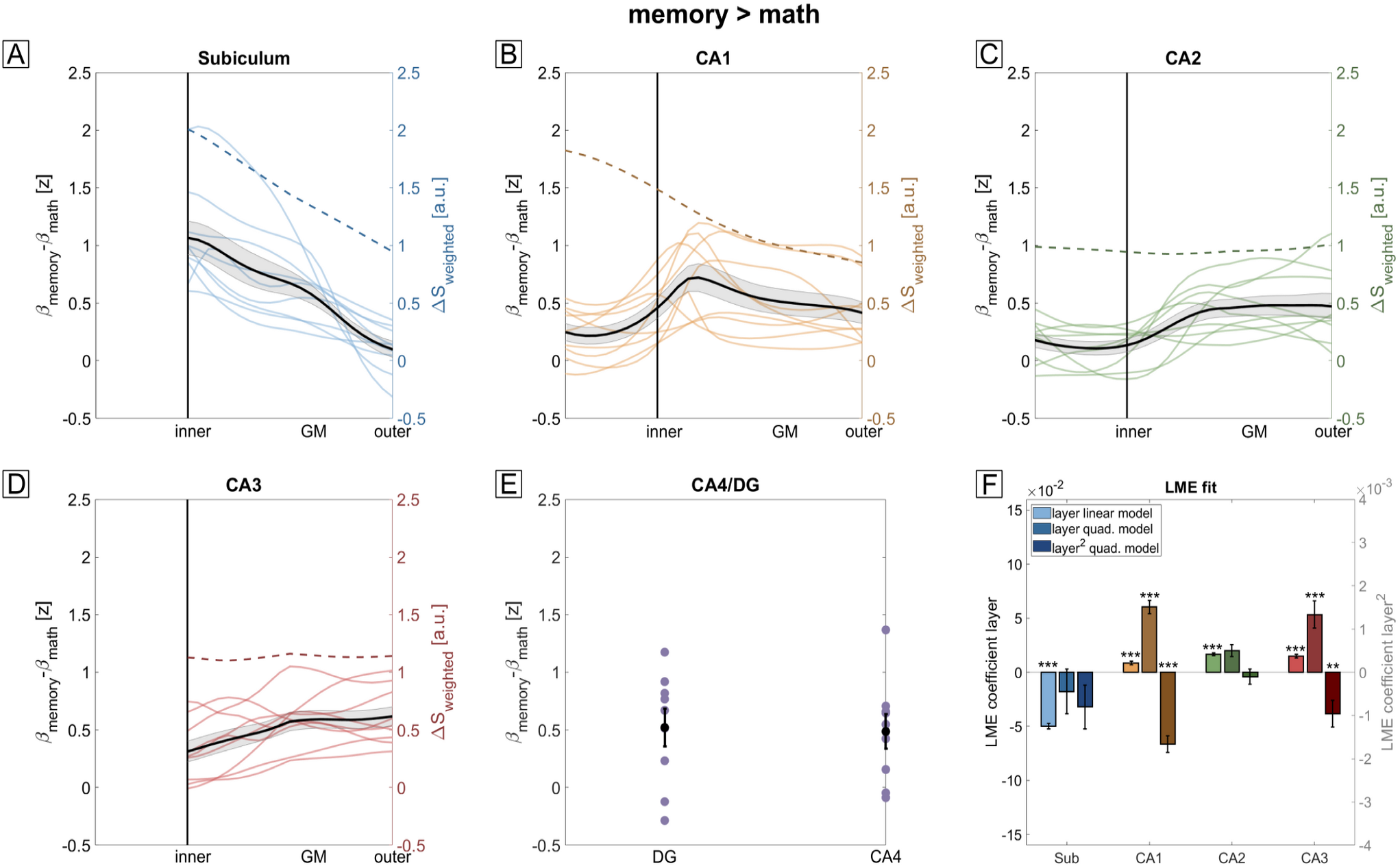
Laminar fMRI profiles of autobiographical memory. **A-E** Laminar fMRI responses for all subfields during the memory > math contrast. Subject-averaged profiles are shown as black overlays (shaded area corresponds to SEM). The vascular-weighted signal change during breath-hold is plotted as dashed line. In case of CA4 and the DG, only two bins were defined since layers are less clearly defined. **F** LME model comparisons using approximated F-tests show a significant effect of depth for all subfields. Here, ‘layer quad. model’ and ‘layer^2^ quad. model’ refer to the linear and quadratic term in the second-order LME model, respectively. In CA1 and CA3, a quadratic model fitted the data significantly better than a linear model. Note that the peak at inner layers of CA1 (B) cannot be explained by a venous bias as this would predict a monotonic increase toward St. lacunosum-moleculare (dashed line).

As autobiographical memory recall exhibits a temporal dynamic^30^, we investigated whether possible laminar differences between memory construction vs. memory elaboration exist. Fig. 3 shows the laminar profiles of this contrast (pre > post). Again, LME analysis showed a highly significant linear main effect of depth (F(1,260) > 104, p_FDR_ < .001). In case of CA3, LME analysis showed a significant quadratic effect of depth (F(1,178) = 37, p_FDR_ = 3e-8) suggesting that recurrent connections within CA3 play a more important role during memory construction than elaboration.

**Fig. 3:**
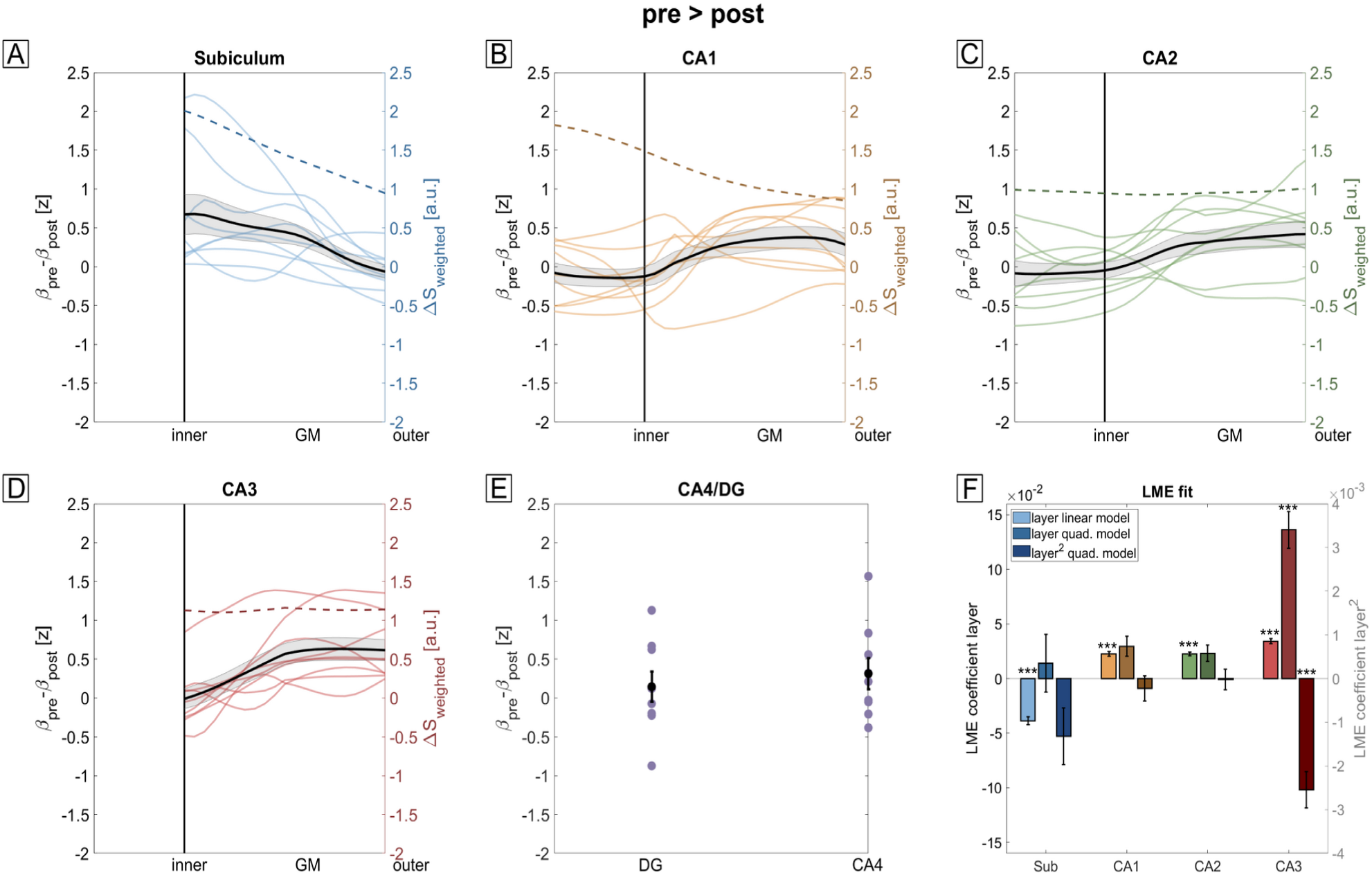
Laminar fMRI profiles of construction vs. elaboration phases. **A-E** Laminar fMRI responses for all subfields during the pre > post contrast. Subject-averaged profiles are shown as black overlays (shaded area corresponds to SEM). The vascular-weighted signal change during breath-hold is plotted as dashed line. **F** LME comparisons using approximated F-tests show a significant effect of depth for all subfields. In CA3, a quadratic model fitted the data significantly better than the linear model. The profiles in CA1 (B) show a significantly different shape compared to the memory > math contrast (Fig. 2B).

We compared the two functional contrasts (memory > math vs. pre > post) by performing a LME analysis for each subfield (except CA4/DG) with depth, contrast and their interaction as main effects (see section 2.2.4 for details). All subfields show a significant difference between contrasts (F(1,528) > 18, p_FDR_ < .001 for all subfields) and a significant interaction between depth and contrast (F(1,528) > 4, p_FDR_ < .05 for all subfields). The strongest effects of contrast and interaction are seen for CA1 (F(1,528) = 101,p_FDR_ = 3e-21 for contrast, F(1,528) = 20.5, p_FDR_ = 3e-5 for the interaction) where the laminar profiles in Fig. 3B do not exhibit a peak at inner layers as shown in Fig. 2B but rather a maximum in mid to outer layers. Importantly, this indicates that laminar profiles depend on the exact task demands (i.e. contrasts) rather than reflecting unspecific differences of apparent overall recruitment that may be confounded by vascular (or other) factors.

In summary, these results show that we can obtain consistent laminar fMRI responses from all subfields of the human hippocampus using GRE-BOLD. Specifically, the subiculum shows on average the strongest BOLD activity in the memory > math condition, in line with Leelaarporn et al.^29^. Furthermore, CA1 shows laminar responses that differ strongly between contrasts and are not explainable by a venous bias alone. These results can serve as a benchmark for future laminar fMRI studies of the human hippocampus at UHF.

#### 3.2.1 Effects of venous masking

We were interested in whether using a mask of venous vessels as obtained from the SWI image (see section 3.1) would significantly reduce the venous bias in high-resolution GRE-BOLD fMRI. Fig. 4 displays the difference between laminar profiles without and with venous masking for both contrasts. To assess whether venous masking has an effect on the laminar profiles we performed a LME analysis similar to that used in the previous section (see section 2.2.4). For the memory > math contrast (Fig. 4A), LME analysis revealed a significant main effect of masking only for the subiculum (F(1,366) = 9.76, p_FDR_ = 0.005). The effect is the strongest at the inner surface as predicted by the hyperemia experiment (Fig. 1H). In case of the pre > post contrast (Fig. 4B), masking had a weak effect on CA2 (F(1,528) = 5.47, p_FDR_ = 0.049). CA3 showed a significant interaction between depth and masking (F(1,366) = 9.05, p_FDR_ = 0.007) but no main effect of masking (F(1,366) = 4.36, p_FDR_ = 0.06).

**Fig. 4:**
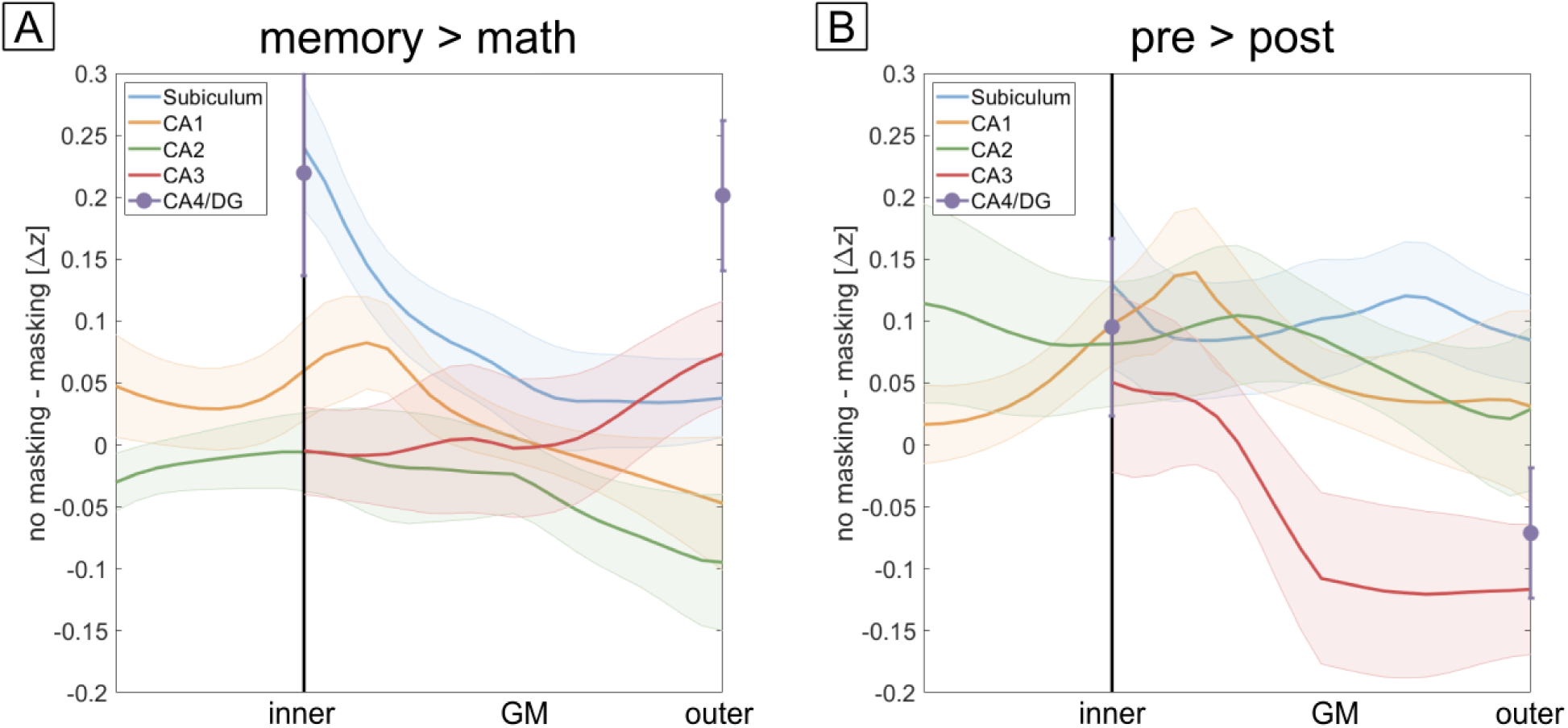
Effect of vein masking. **A** Difference in functional contrast (memory > math) when performing vein masking. Only the subiculum shows a statistically significant effect of masking and the effect is the strongest at the inner surface where the bias is the strongest (see Fig. 1H). **B** Same as in **A** but for pre > post. CA2 shows a weak effect of masking (p = 0.049) and CA3 shows a significant interaction between depth and masking but no main effect of masking.

In summary, these results indicate that vein masking has a noticeable effect on the subiculum which is shown to be most affected by the venous bias. Furthermore, venous masking has a stronger effect during the memory > math contrast compared to the pre > post contrast.

### 3.3 MRI metrics

In addition to the laminar fMRI results, the experiments performed in this study allowed us to obtain novel insights into depth-dependent MRI metrics of the human hippocampus relevant for the design and evaluation of other methods than GRE-BOLD.

#### 3.3.1 T_2_*

By fitting a mono-exponential decay function to the multi-echo data during normocapnia (sections 3.1, 2.1.3), we were able to obtain baseline T_2_* relaxation times for each subfield as a function of cortical depth. Fig. 5A-D shows the fitted T_2_* on the unfolded inner (A) and outer (B) surface as well as on the folded outer surface (C). Aggregating vertices results in the depth-dependent plots shown in Fig. 5D. Interestingly, the subiculum exhibits a comparably short T_2_* relative to the remaining subfields. This can be explained by a high myelin content in the subiculum as noted with histology^71^.

**Fig. 5:**
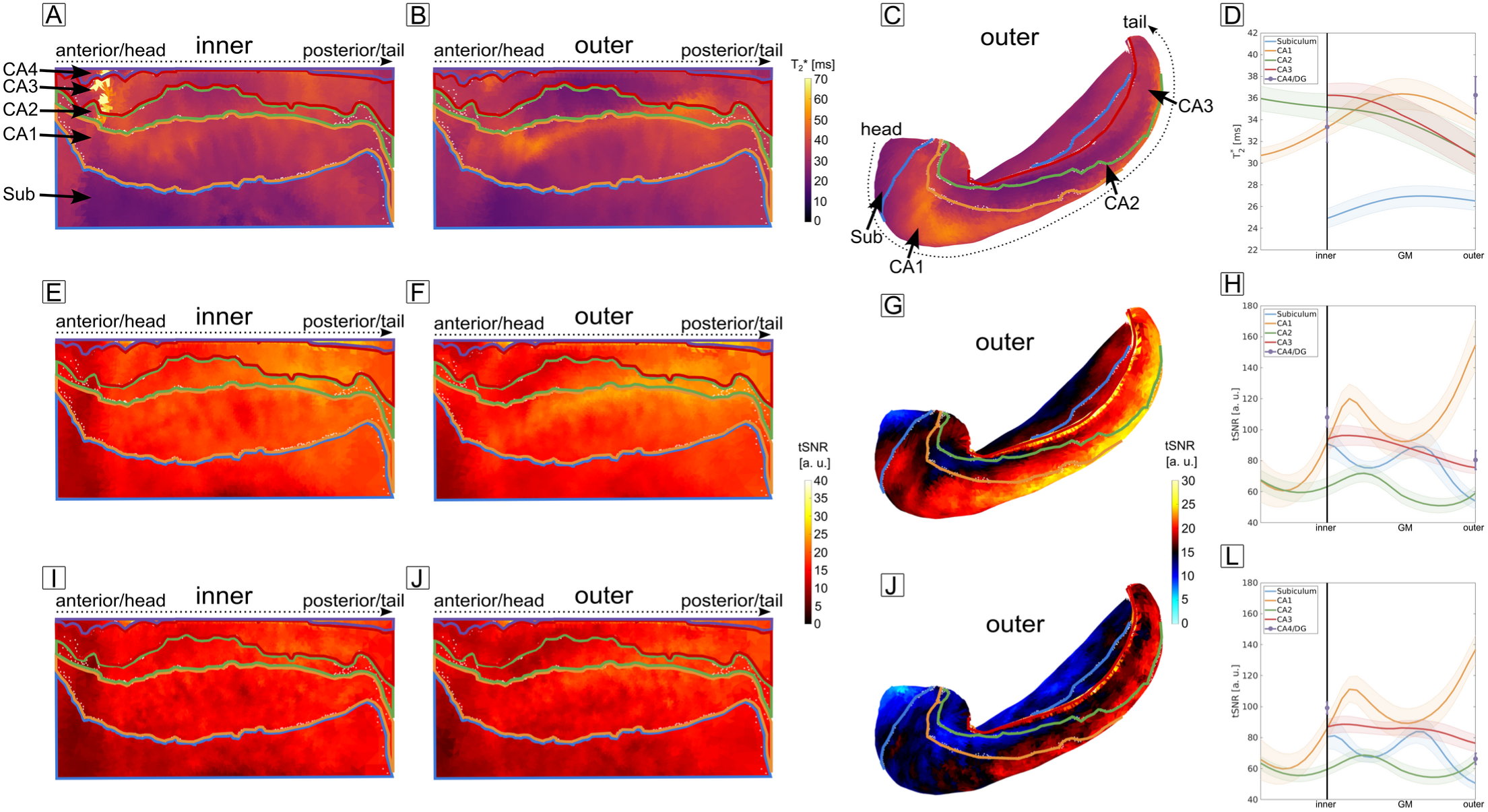
MRI metrics. **A-B** 2* relaxation time obtained during the breath-hold experiment and projected onto the unfolded hippocampal inner (**A**) and outer (**B**) surface. **C** Same as in **B** but shown on the folded hippocampal surface. The structure of long T2* at the demarcation between CA1 an CA2 is likely due to partial voluming with CSF. **D** Laminar profiles of T2* for all subfields. The subiculum exhibits a comparably short T2* relative to the remaining subfields. **E-F** tSNR maps obtained during the AM experiment without venous vessel masking projected onto the unfolded hippocampal surfaces. **G** Same as in **F** but shown on the folded hippocampus. **H** Laminar profiles of tSNR. The high tSNR at the outer layer of CA1 can be explained by partial voluming with CSF. **I-J** Same plot as shown in **E-H** but for the case of venous vessel masking. Although an overall reduction of tSNR is present, this reduction is negligible when averaging over vertices (**L).**

The outer surface as shown in Fig. 5B,C exhibits a noteworthy area of long T_2_* at the demarcation between CA1 and CA2. The most likely cause for this is a partial voluming effect with CSF in the proximity of the hippocampus (cf. supplementary Figure S4).

#### 3.3.2 tSNR

As a measure of overall signal stability, temporal SNR is often reported. In this study, we obtained tSNR on the voxel-level (tSNR of two subjects scanned twice on different days shown in supplementary Fig. S8) and as a function of cortical depth by calculating the temporal mean and standard deviation over volumes corresponding to the control condition of the AM experiment. In Fig. 5E-G, the tSNR is shown as a projection on the unfolded inner and outer hippocampal surface, as well as the folded outer surface. Here, no masking for venous vessels was performed. The same structure of long T_2_* is now seen as high tSNR at the demarcation between CA1 and CA2, resulting in an overall high tSNR at the outer surface of CA1 as shown in Fig. 5H. This result supports the notion that partial voluming with CSF is the most likely cause.

Masking venous vessels (Fig. 5I-L) led to an overall reduction in tSNR as the masking process also excluded GM vertices which are partial voluming with veins. However, when averaging over vertices to obtain layer profiles (Fig. 5L), the reduction in tSNR is negligible (i.e. comparing Fig. 5H with 5L).

#### 3.3.3 Physiological noise

An important question to answer is whether the noise in the laminar profiles is dominated by physiological noise, i.e. whether averaging does not further improve the tSNR ^61^. To investigate the laminar noise characteristics, we performed a Weisskoff test for each subfield^65^ (see section 2.2.3.2). Fig. 6A-E shows the subject-averaged temporal noise as a function of cortical depth and amount of averaged vertices 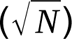. For all subfields, the noise does not vary considerably with averaging when approaching the maximum number of available vertices. In particular, the physiological noise regime is reached when approximately 2300, 5700, 1100, 1300, and 3400 vertices are averaged for the subiculum and CA1-4, respectively. The number of vertices is well below the total number of vertices available for averaging in each subfield highlighting that in our current experimental setup, we can safely assume to be in the physiological noise dominated regime.

**Fig. 6:**
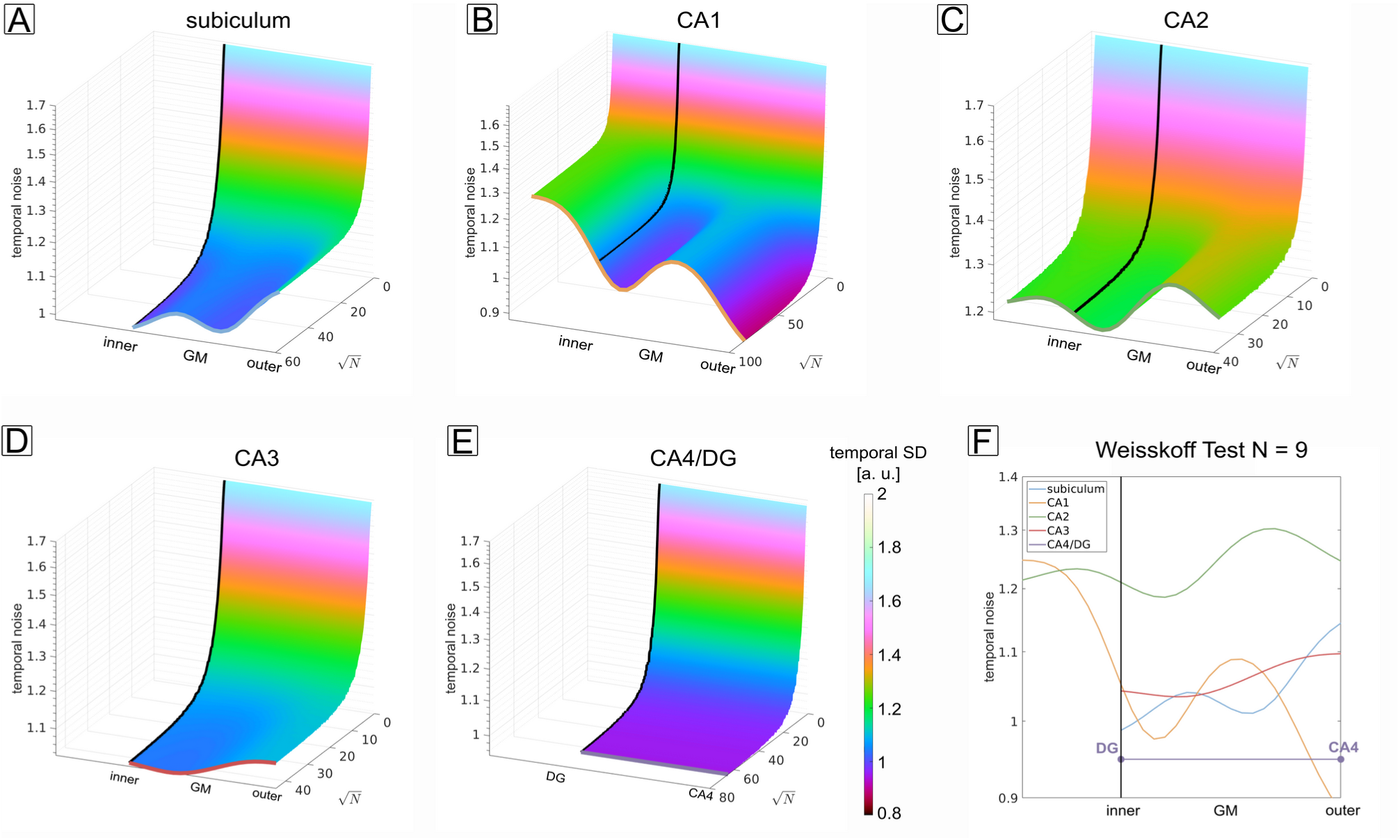
Weisskoff test and physiological noise. **A-E** Subject-averaged temporal SD as a function of cortical depth and the amount of averaged vertices for all subfields. When approaching the maximum number of available vertices, the noise does not decrease considerably with averaging showing that the remaining patterns stem from physiological noise. F Noise profiles in the physiological noise dominated regime, i.e. taking the profiles when all vertices were averaged (colored lines in A-E).

The noise profiles in the physiological noise regime (Fig. 6F) exhibit a subfield-specific laminar structure. Koopmans et al.^66^ argued that the physiological noise profiles follow the vascular density and hence give estimates of blood volume^72^. We also sampled an image of an India Ink stain (supplementary figure S9C, adapted from Duvernoy et al. ^48^), to compare profiles of the vascular density with the obtained physiological noise profiles. Results can be found in the supplementary material (Fig. S9).

## 4. Discussion

In this study, we show that is it possible to image the human hippocampus at a precision high enough to robustly sample BOLD responses and other physiological and MR-related parameters as a function of cortical depth in each individual hippocampal subfield.

The results of the breath-hold experiment indicate that among all subfields, the subiculum, closely followed by CA1, has the strongest venous bias, blurring activity toward inner layers (Fig. 1H). By mapping venous vasculature (Fig. 1A-G), we could show that the bias is driven by a heterogenous distribution of large venous vessels across layers of the hippocampus. This result was to be expected: The GRE-BOLD contrast, being driven by venous vessels, scales with vessel size^20^ and follows the underlying vascular architecture^17,19^. In light of hippocampus-specific features, a recent study by Haast et al.^8^ additionally showed that the subiculum is proximal to arterial macrovessels. Therefore, we advise interpreting strong laminar fMRI responses in inner layers of the subiculum, but also CA1, with caution. Although lower in magnitude, CA2/3 show a weak bias toward the outer surface, hence any laminar trend toward outer layers in these subfields needs to be interpreted carefully.

The anatomical location of the hippocampus renders it prone to artifacts associated with B_0_-inhomogeneity. Although the acquisition of phase stabilization echoes strongly improved image quality and stability (supplementary figure S2), performing a breath-hold experiment by itself inevitably amplifies B_0_-inhomogeneity, especially at longer TE. We tried to reduce these artifacts by down-weighting longer echoes prior to echo averaging. In this regard, attempts to reduce venous bias by performing a breath-hold calibration scan at typical echo times when using an EPI readout might be less successful in the hippocampus compared to the neocortex^73,74^.

By acquiring 3D-EPI images at 0.9 mm isotropic resolution, we could obtain GRE-BOLD responses as a function of cortical depth during an autobiographical memory paradigm (Fig. 2, Fig. 3). Comparing these results with those of the hyperemia experiment (Fig. 1), CA1 shows a clearly different behavior than what would be predicted by a venous bias alone. If the venous vasculature distribution was the sole driver of these effects, a peak in St. lacunosum-moleculare of CA1 would be expected in both contrasts (dashed orange line in Fig. 2B and Fig. 3B). However, this is not what we see. In the memory > math contrast (Fig. 2B), the activity increases from outer to inner layers and peaks at the inner layer (around the putative transition between St. pyramidale and St. radiatum) before leveling off when approaching the most inner layers St. lacunosum-moleculare. This peak suggests a predominant relevance of inputs via the trisynaptic pathway, i.e. from CA3, rather than of direct EC inputs during autobiographical memory retrieval. In the pre > post contrast, the profiles of CA1 (Fig. 3B) show a significantly different behavior compared to the memory > math contrast. Here, no difference between construction and elaboration phase exists in the innermost layers (putatively St. lacunosum-moleculare). Instead, the contrast increases from inner to outer layers, suggesting that trisynaptic path inputs from CA3 are more relevant during memory construction than elaboration.

Laminar differences of hippocampal BOLD responses may depend on multiple different factors, which should be disentangled in the future and the current results can serve as a baseline the community can refer to. Some factors to consider would be differences between the laminar distribution of “inputs”, i.e. synaptic inputs (which may arrive at all layers) vs. “outputs”, i.e. action potentials (which are generated perisomatically, i.e. St. pyramidale) or differences between different types of synaptic inputs (largely entorhinal inputs vs. perforant path inputs). These could be tested via functional connectivity analyses, e.g. via PPI^75^. In addition, differences in laminar distribution of inhibitory interneurons might also play a role. The latter could be tested pharmacologically by combining laminar fMRI with an intervention such as application of benzodiazepines, (which are GABA-A receptor agonists) or by studying epilepsy patients with altered function in specific populations of interneurons.

Accompanying the laminar fMRI results, our study also provides insights into MRI and physiological parameters at the mesoscale. Our T_2_* profiles are in line with previous reported T_2_* values obtained at lower spatial resolution at 7 T^76^ and the ability to obtain depth-dependent profiles adds a new dimension to investigate MRI parameter differences between healthy controls and patients^76^. From an fMRI acquisition perspective, our results can help to guide researchers in selecting echo times optimal for their specific study. In particular, Fig. 5D shows that a longer TE than used in this study would be theoretically more optimal in terms of maximizing BOLD sensitivity. However, the breath-hold results indicate what artifact level can be expected at longer TE. Because we could obtain robust BOLD responses at a TE of 22 ms, we advise scanning at a shorter than theoretically optimal TE to improve temporal efficiency and signal stability.

The obtained tSNR maps (Fig. 5, supplementary figure S8) and profiles indicate that we could obtain robust laminar fMRI responses using 3D-EPI. Excluding vertices shown to be likely co-located with veins reduced tSNR but the effect masking has on the tSNR of aggregated laminar profiles is negligible and it reduces signal changes at surfaces known to be prone to venous bias (Fig. 4). However, it must be noted that the measures to reduce venous bias used in this study can only serve as a first order correction as the BOLD effect of large veins affects all depths of the parenchyma^77,78^ and the convolutional effect of draining veins^79^ cannot be accounted for with masking. Hence, other fMRI contrasts like SE-BOLD^80^ or non-BOLD alternatives like VASO^12,15,81^ are needed to be developed further to obtain laminar fMRI responses with an as low as possible effect of large vessels and our results can serve as a benchmark to compare these contrasts to. Alternatively, other post-processing methods like phase regression^82^ could also be employed to reduce venous bias although we were not able to reliably reduce effects of large veins using phase regression, most likely due to differences in SNR between the hippocampus and the neocortex where this method has previously been used (data not shown). In addition to the methods mentioned here, the design of the neuroscientific experiment can also help to reduce effects of blood drainage^83^. In this study, we contrasted two phases that putatively both involve the hippocampus (construction vs. elaboration phase). Alternatively, modulating the strength of different hippocampal inputs, similar to work performed in sensory areas^84^, might be a possibility to control the vascular bias in GRE-BOLD.

The noise curves in Fig. 6 all reach a plateau, showing that when aggregating over vertices, physiological noise is the dominant noise source. This is in line with previous studies^66,67^, indicating that the effect of thermal noise can be neglected. However, given the hippocampus’ susceptibility to signal fluctuations driven by physiology, some measures of noise reduction need to be applied to increase data reproducibility, especially in inferior brain regions. We demonstrated this by assessing model residuals before and after using acompCor (Supplementary figure S5) showing that modeling physiological noise sources in the GLM is advisable as it increases the model’s predictability. Future work could include a more systematic evaluation of acompCor and other methods, e.g. RETROICOR ^85^. Our own attempts^86^ to compare acompCor and RETROICOR show a superior performance of acompCor, possibly due to a more robust estimation of physiological noise sources when using acompCor.

In conclusion, sampling BOLD responses as a function of cortical depth makes investigation of the underlying functional microcircuits of the human hippocampus possible, and has potential for non-invasively investigating the directionality of the projections to and from other brain regions. Here, we present relevant insights on the vascular bias of laminar fMRI of the human hippocampus which will help the community to interpret hippocampal responses when utilizing the most widely used fMRI contrast, GRE-BOLD. Finally, the presented AM results and methods will help researchers to test new hypotheses on a laminar level and to validate other fMRI contrasts to improve the specificity of laminar fMRI of the human hippocampus.

## 5. Acknowledgements

The authors thank Roy Haast for his helpful comments on hippocampal surface generation and for providing code for the online app. The authors would also like to thank the Khan Lab for their support in using hippunfold. The authors thank Rüdiger Stirnberg and the team at DZNE Bonn for providing the 3D-EPI sequence. The research leading to these results has received funding from MERCUR grant number Ko-2021-0010. The MAGNETOM Terra used in the study was funded by the Deutsche Forschungsgemeinschaft (DFG, German Research Foundation), grant number 432647511.

