## Supplementary Material for "Characterizing BOLD activation patterns in the human hippocampus with laminar fMRI"

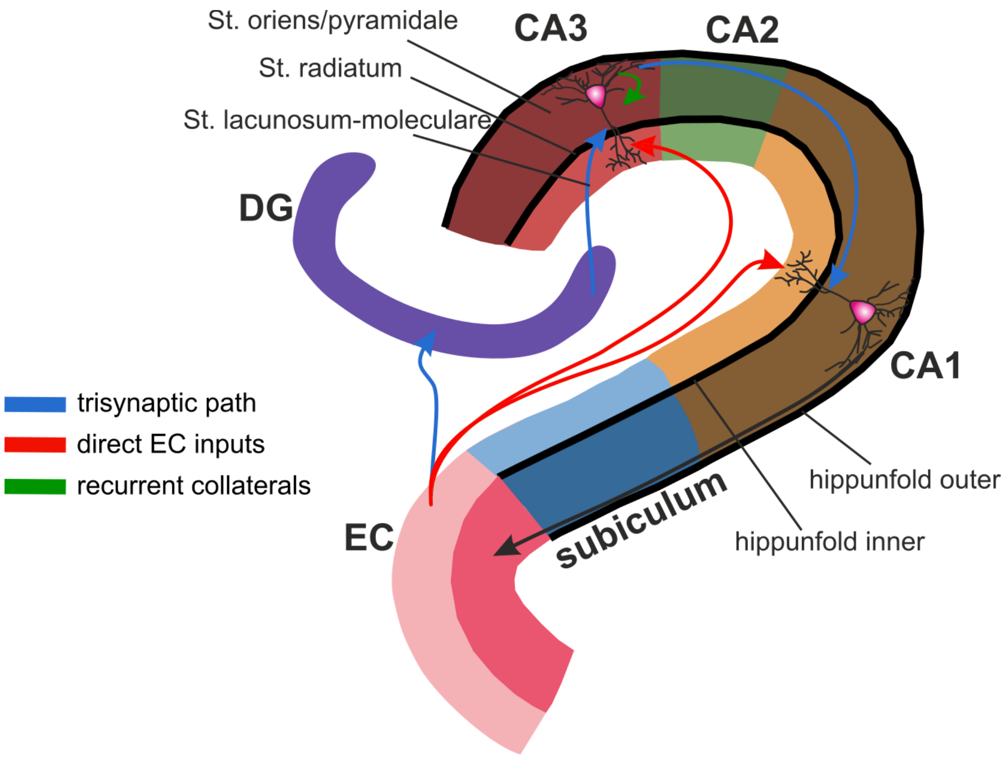


**Fig. S1: Neuronal pathways of the hippocampus.** Schematic depiction of the neuronal pathways of the hippocampus. Inner layers are in lighter colors, outer layers in darker colors. The surfaces as given by hippunfold[1] are shown for reference.

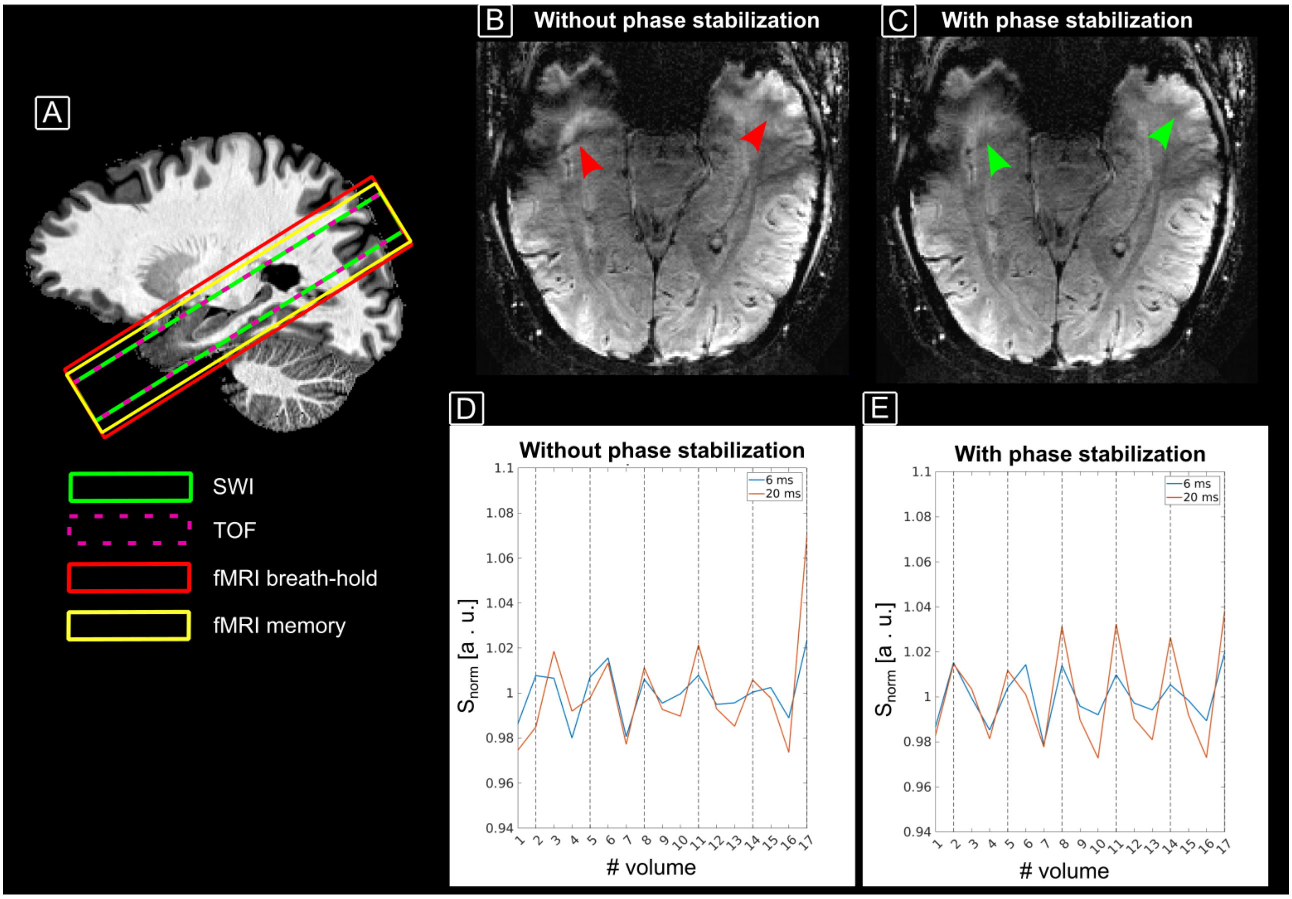


**Fig. S2: FOV placement and breath-hold signal stability. A** FOV placement of each MRI contrast with respect to the anatomical reference. **B-C** Images acquired during the inhalation period (transition phase) of the last echo (TE = 21 ms) without (**B**) and with (**C**) phase stabilization. $B_0$-induced artifacts (red arrows in **B**) are reduced with phase stabilization (green arrows in **C**). **D-E** Time series of the echo-normalized signal obtained from the inner surface of the hippocampus for TE = 6 ms (blue lines) and 20 ms (red lines). The dashed vertical lines correspond to an 'active' volume. With phase stabilization (**E**), the signal change is more consistent, and coincides well with the paradigm, especially at longer echoes.

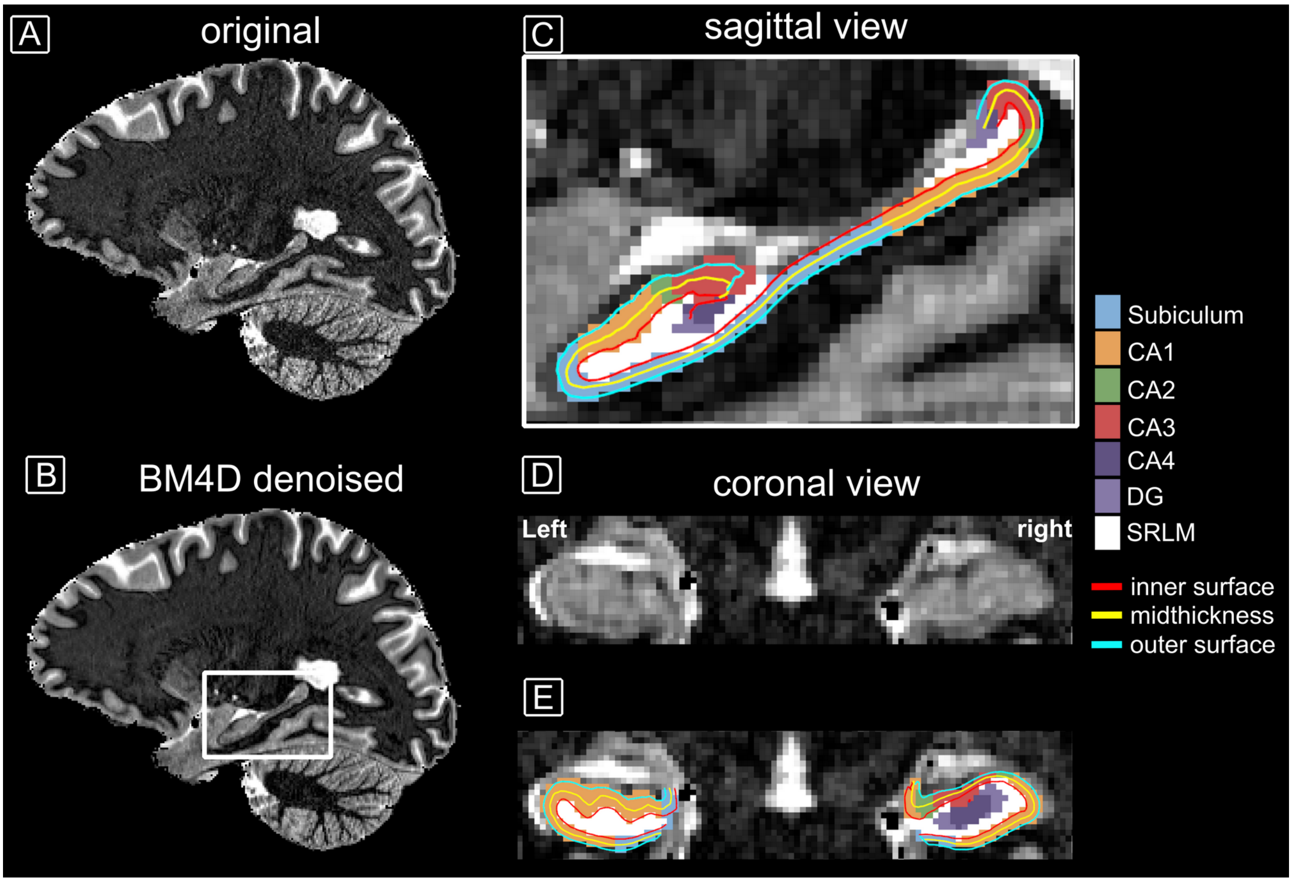


**Fig. S3: Denoising performance and hippocampal segmentation. A** Original $T_1$-weighted image (first inversion time of MP2RAGE, i.e. INV1). **B** INV1 after denoising using BM4D. Salt and pepper noise is strongly suppressed while fine-scale details are preserved. **C-E** Segmentations and surfaces generated by hippunfold when using INV1 as a substitute for a $T_2$-weighted image.

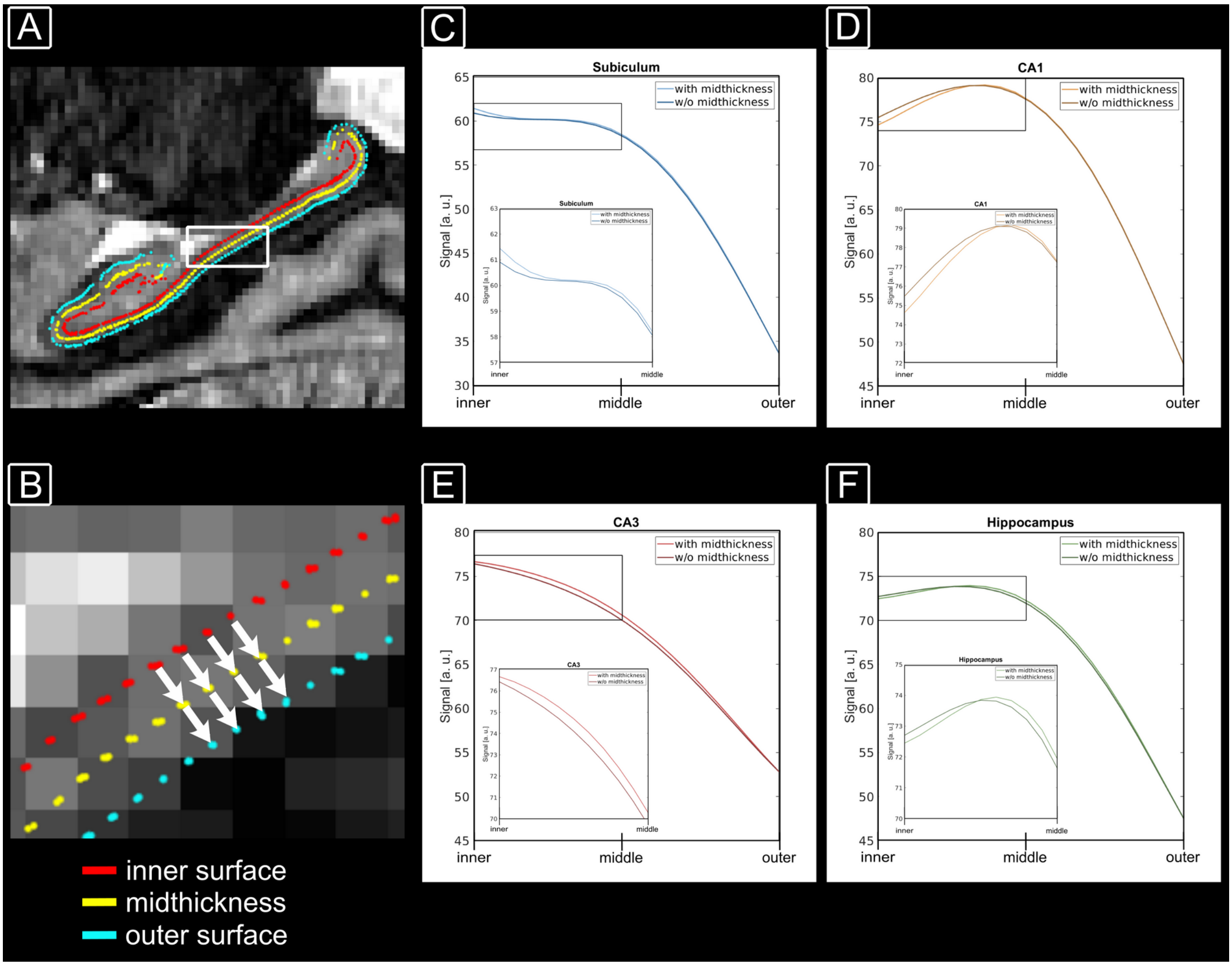


**Fig. S4: Hippocampus laminar sampling strategy. A** Surfaces obtained with hippunfold. **B** Zoomed section in **A**. Each vertex on the inner surface has a counterpart on the outer surface. For these vertex pairs, the signal is sampled, taking the middle surface as an anchor point into account (white arrows in **B**). This strategy is similar to that presented by Koopmans et al.[3]. **C-F** Signal sampled from the INV1 image of the MP2RAGE (**A-B**) for example subregions and for the entire hippocampus with and without taking the midthickness surface into account. Zoomed sections are shown as inlays. The sampling strategy essentially obeys the equivolume principle[4], but similar to the findings of Waehnert et al.[5] the profiles do not strongly differ from those obtained with a pure equidistant sampling given the still relatively coarse resolution of typical laminar fMRI experiments.

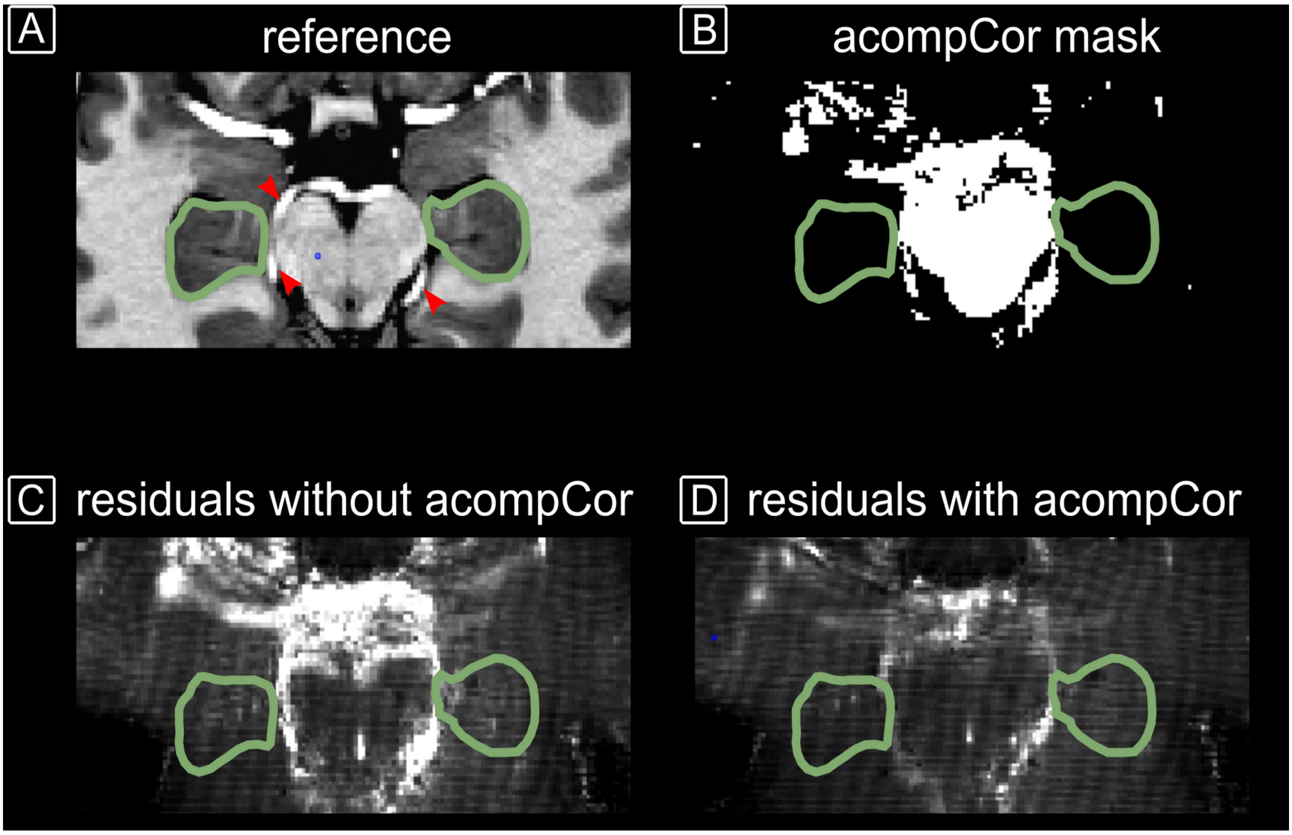


**Fig. S5: Physiological noise correction strategy.** Anatomical reference of an exemplary subject. The hippocampi are highlighted as green ROIs. Note the large arteries (red arrows) close to the hippocampus. **B** Mask used for acompCor regression. It contains the manually drawn brainstem white matter mask and a mask of high residuals as given by the first GLM. **C** Residuals of the first GLM. **D** Residuals after the second GLM using approx. 13 acompCor regressors. The residuals within the hippocampus reduced from 195 ± 76 to 170 ± 36.

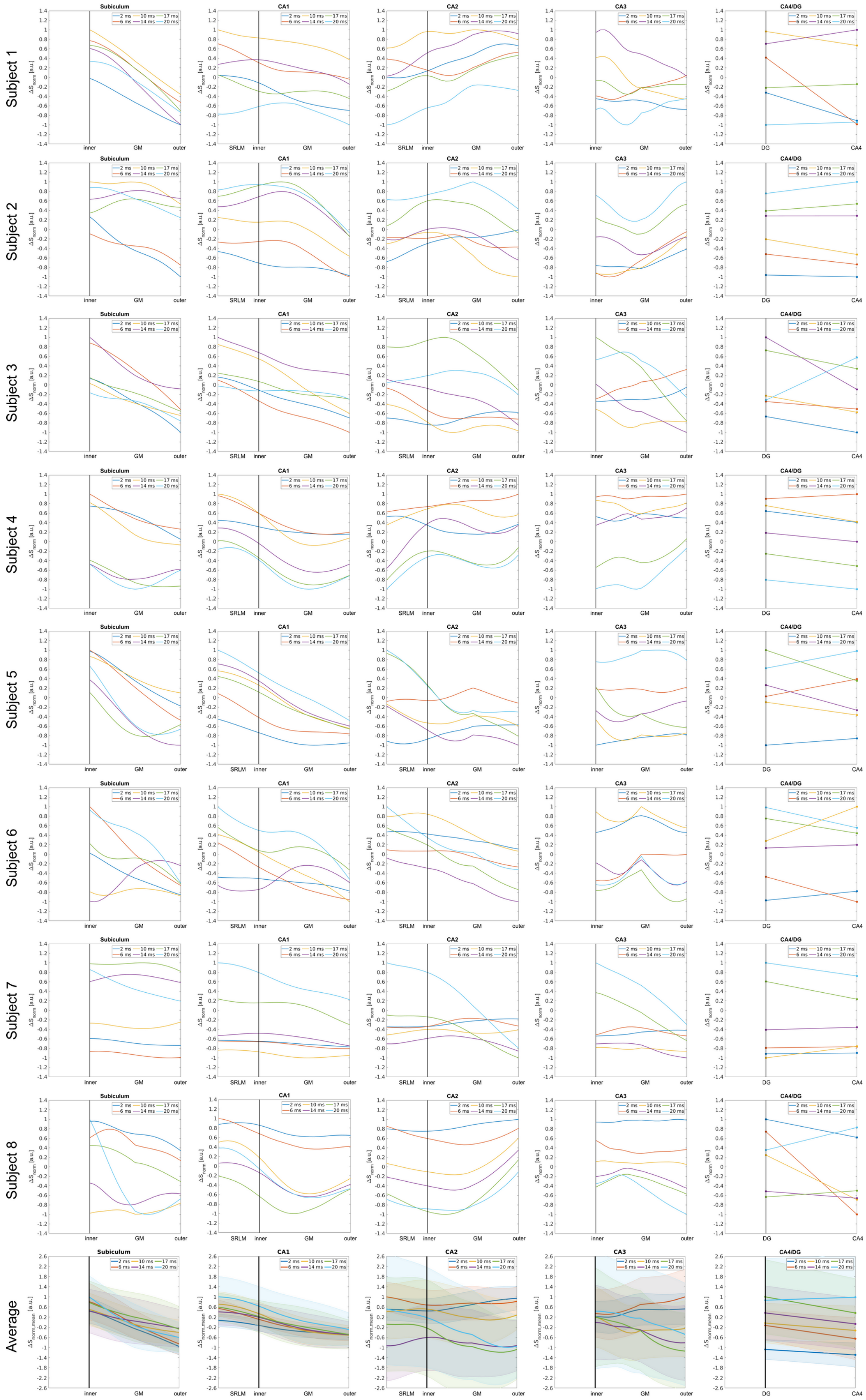


**Fig. S6: Single subject and subject-averaged multi-echo breath-hold results.** Profiles are shown for each subject and subfield. The shaded area in the average plots (bottom row) is the SEM.

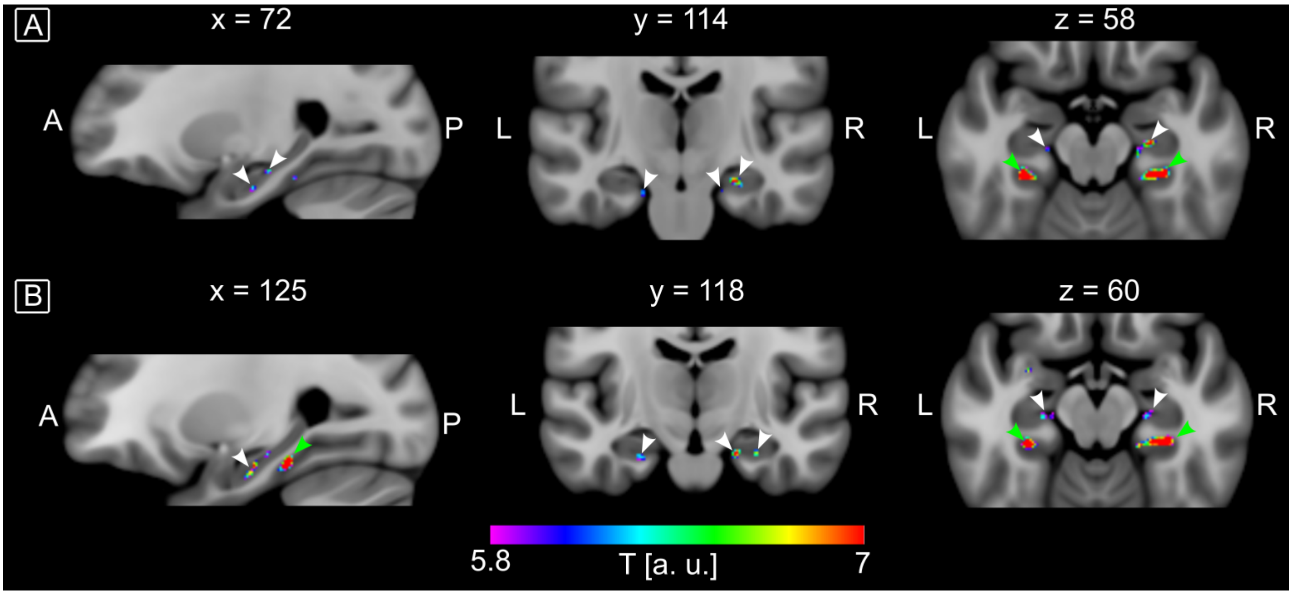


**Fig. S7: Voxel level group results.** Maps of pseudo t-values at the group level as computed with the SnPM toolbox ($p < .05$, FWE-corrected, coordinates given in MNI space). The maps show the expected activation patterns for the hippocampus (white arrows, see also Fig. 1 in Leelaarporn et al.[2]) and the parahippocampal gyri (green arrows) when performing the autobiographical memory paradigm.

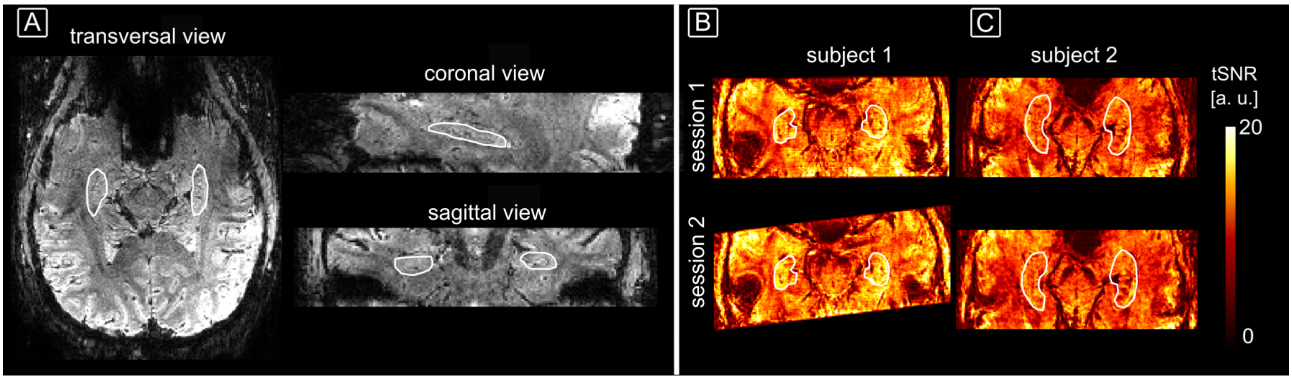


**Fig. S8: Autobiographical memory experiment: image quality. A** Example single volume of the first subject (in native space). The hippocampus is highlighted as white ROI. **B-C** tSNR maps of both sessions for subject 1 (**B**) and subject 2 (**C**) (in anatomical space). The tSNR in the hippocampus averaged over both hemispheres (white ROIs) was similar between sessions (Subject 1: 11.3 ± 3.2 for session 1, 11.4 ± 3.2 for session 2; Subject 2: 8.4 ± 2.5 and 8.6 ± 2.6, respectively).

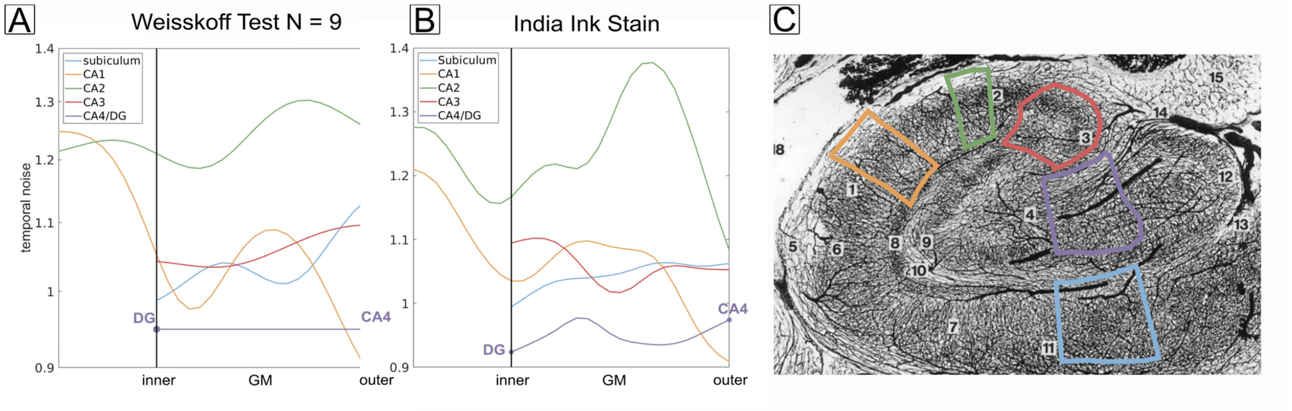


**Fig. S9: Comparison between profiles of physiological noise and microvascular density. A** Noise profiles in the physiological noise dominated regime (same as Fig. 6F of the main manuscript). We sampled the pixel intensity in the colored regions highlighted in **C** by adapting the approach as described in (**https://layerfmri.com/2018/07/19/2dlayers/#more-950**). The regions mark parts of the different subfields (image adapted from Duvernoy et al.[6]). **B** Obtained profiles smoothed to match the effective resolution of the main experiment. The profiles show a strong resemblance to those obtained from the Weisskoff test (A) indicating that the variation of physiological noise profiles could be caused by the underlying blood volume distribution.
